# Museum genomics reveals temporal genetic stasis and global genetic diversity in *Arabidopsis thaliana*

**DOI:** 10.1101/2025.02.06.636844

**Authors:** Lua Lopez, Patricia L. M. Lang, Stephanie Marciniak, Logan Kistler, Sergio M. Latorre, Asnake Haile, Eleanna Vasquez Cerda, Diana Gamba, Yuxing Xu, Patrick Woods, Mistire Yifru, Jeffrey Kerby, John K. McKay, Christopher G. Oakley, Jon Ågren, Tigist Wondimu, Collins Bulafu, George H. Perry, Hernán A. Burbano, Jesse R. Lasky

## Abstract

Global patterns of population genetic variation through time offer a window into evolutionary processes that maintain diversity. Over time, lineages may expand or contract their distribution, causing turnover in population genetic composition. At individual loci, migration, drift, and selection (among other processes) may affect allele frequencies. Museum specimens of widely distributed species offer a unique window into the genetics of understudied populations and changes over time. Here, we sequenced genomes of 130 herbarium specimens and 91 new field collections of *Arabidopsis thaliana* and combined these with published genomes. We sought a broader view of genomic diversity across the species, and to test if population genomic composition is changing through time. We documented extensive and previously uncharacterized diversity in a range of populations in Africa, populations that are under threat from anthropogenic climate change. Through time, we did not find dramatic changes in genomic composition of populations. Instead, we found a pattern of genetic change every 100 years of the same magnitude seen when comparing Eurasian populations that are 185 km apart, potentially due to a combination of drift and changing selection. We found only mixed signals of polygenic adaptation at phenology and physiology QTL. We did find that genes conserved across eudicots show altered levels of directional allele frequency change, potentially due to variable purifying and background selection. Our study highlights how museum specimens can reveal new dimensions of population diversity and show how wild populations are evolving in recent history.

## Introduction

The genomic composition and diversity of species across their geographic range can reveal the relative importance of processes such as demography, drift, and spatially varying selection. The evolutionary history of a species is reflected in its present-day patterns of relatedness across the genome, and these patterns will change in the future as demography and environments change (Fulgione & Hancock, 2018; C.-R. Lee et al., 2017; Olalde et al., 2018; Rhoné et al., 2020). For example, regions that differ in their history of colonization may show distinct patterns of diversity and genetic turnover through space (Lasky et al., 2024; Petkova et al., 2016). Furthermore, knowledge of geographic patterns in diversity and population structure can be used to optimize sampling designs for studies of genetic variation in phenotype (Fulgione & Hancock, 2018).

While biologists have often studied geographic patterns in genotype and phenotype, their study of the temporal genomic dynamics has developed only more recently, largely due to a previous lack of data. As global environments are changing, species range-wide study of temporal dynamics in population genomics could reveal corresponding evolutionary changes. However, the breadth of sampling required through space and time is a logistical hurdle. Recent studies have used ancient human sequences to demonstrate dramatic turnovers in population ancestry in some regions, as new lineages replaced older ones (Allentoft et al., 2024; Kennett et al., 2022; LaPolice et al., 2024; Olalde et al., 2018), or consistent geographic population structure through time (Antonio et al., 2024). For non-human species, long-term field studies have offered powerful windows into temporal dynamics (Bergland et al., 2014; Lynch et al., 2024; Troth et al., 2018). Global natural history collections made over the last couple of centuries are an underutilized but incredibly valuable resource for biology, holding a wealth of information on genotype and phenotype (Burbano & Gutaker, 2023; Lopez et al., 2020). For example, researchers have used museum specimens to show demographic turnover in a plant species where there was dramatic expansion of tetraploids across parts of Europe formerly inhabited only by diploids (Rosche et al., 2024). Recent population studies have also used sequences of museum specimens to identify changes in genetic diversity over time (Bi et al., 2019; Gauthier et al., 2020) and the history of spread of plant pathogens (Yoshida et al., 2013). Furthermore, museum collections often hold samples from a wide geographic area, allowing greater spatial coverage in evolutionary studies.

*Arabidopsis thaliana,* the model plant (hereafter referred to as “Arabidopsis”), is a useful system for studying drivers of geographic and temporal turnover in genotype. Arabidopsis is an annual, largely self-fertilizing plant species with a broad native distribution in Eurasia and Africa (Fulgione & Hancock, 2018; Yim et al., 2023). Populations of Arabidopsis occur across a wide range of climatic regions and different environments (e.g., from the Mediterranean coast to alpine systems) (Ågren & Schemske, 2012; Elfarargi et al., 2023; Gamba et al., 2024; Montesinos et al., 2009; Yim et al., 2023). Global studies on *A. thaliana* have revealed high levels of genetic and phenotypic diversity among populations (Koornneef et al., 2004; Mitchell-Olds, 2001) and this species has become a model for eco-evolutionary studies (Takou et al., 2019; Turner et al., 2020; Vasseur et al., 2018; Wuest & Niklaus, 2018).

Along with life history variation across contrasting environments, *A. thaliana* displays substantial large-scale population structure (Beck et al., 2008; Lasky et al., 2012; Schmid et al., 2006; Sharbel et al., 2000). The largest genetic divisions within this species can be found between most Eurasian genotypes and the so-called relict lineages associated with distinct mountain ranges in Africa, Atlantic islands, and some areas of Mediterranean Europe (Alonso-Blanco et al., 2016; Durvasula et al., 2017; Fulgione & Hancock, 2018; Gamba et al., 2024; C.-R. Lee et al., 2017). However, many such regions, especially in Africa (Yim et al., 2023), have been little-sampled and it is likely there are additional uncharacterized unique lineages. Isolation by distance has also been detected at regional and local scales (Hesen et al., 2024; Lasky et al., 2024; Picó et al., 2008; Tyagi et al., 2016) suggesting that Arabidopsis gene flow is rather limited in space.

Plant populations are often adapted to their local environment (Ågren & Schemske, 2012; Fournier-Level et al., 2011; Hereford, 2009; Lasky et al., 2018; Leimu & Fischer, 2008). Individual genes underlying environmental adaptation in plants have been characterized for a number of these traits. For example, local adaptation has been mapped to natural variation in flowering time and phenology genes (Fulgione et al., 2022; Martínez-Berdeja et al., 2020), genes affecting freezing tolerance (G. Lee et al., 2024; Monroe et al., 2016), trichomes (Arteaga et al., 2022), floral pigmentation (Todesco et al., 2022), and resistance to parasites and pathogens (Bellis et al., 2020; Karasov et al., 2014). However, these loci have been primarily identified and studied in the context of existing contemporary diversity, while loci underlying adaptation to environmental change through time are less well known (but see (Exposito-Alonso, Becker, et al., 2018; Franks et al., 2016; Lang et al., 2024; Troth et al., 2018). Furthermore, across the vast Arabidopsis range major elevational clines in some traits and loci reverse in direction between different mountains, indicating alternative local adaptation strategies (Gamba et al., 2024). This strongly suggests consideration of regional patterns is important to understand spatiotemporal patterns of genomic variation in Arabidopsis.

Environments are always changing, but current anthropogenic changes are especially rapid, and wild organisms are responding in some dramatic ways. Resurrection and long-term experimental studies have shown evolutionary changes in quantitative traits and allele frequencies (Anstett et al., 2024; Hamann et al., 2018; Sekor & Franks, 2018) and studies of museum specimens have shown changes in traits in response to environmental change (DeLeo et al., 2020; MacLean et al., 2018; Ng et al., 2023). In Arabidopsis we documented temporal phenotypic changes in herbarium specimens from the last two centuries, namely in collection date, leaf C:N, and leaf d^15^N (DeLeo et al., 2020). The degree to which these changes are genetic versus plastic, and potential genetic changes are oligogenic versus polygenic, remains unknown. Recently, we used herbarium sequences to show that alleles associated with decreased stomatal density were rising in frequency at multiple loci (Lang et al., 2024; Latorre et al., 2022; Lopez et al., 2022). However, the degree of ancestry turnover in Arabidopsis populations and the potential change in quantitative trait loci (QTL) for ecologically important traits remains unexplored.

Multiple types of selection can influence temporal allele frequency dynamics. Abiotic and biotic environments can undergo a sustained directional shift, such as greenhouse gas-induced warming, causing selection for new traits (Lynch & Lande, 1993). As a result, the underlying allele frequencies may show directional shifts, including sweeps (Hayward & Sella, 2022; Höllinger et al., 2019). Next, new beneficial mutations may rise in frequency, with some sweeping to fixation (Patwa & Wahl, 2008). Other types of selection (e.g. frequency dependence) might cause more complicated temporal dynamics, which we set aside for our current study (Fijarczyk & Babik, 2015; Siewert & Voight, 2017). Here, we focus on directional allele frequency shifts. By identifying genes showing dramatic allele frequency shifts, researchers may gain insight into mechanisms of adaptation to directional shifts in environment. Of course, drift will also cause allele frequency shifts over time, thus it may be necessary to identify loci with strong shifts relative to the genomic background (Akbari et al., 2024).

Despite changing environments, it is likely that many traits and loci do not experience changes in selection and instead are subject to consistent stabilizing or purifying selection. For example, many genes are highly conserved in structure over speciation events and many millions of years (Kachroo et al., 2015; Margoliash, 1963). However, it is important to recognize that purifying selection will still lead to non-random change in frequencies of segregating alleles over time, i.e. decreases in deleterious allele frequency. In simulations, (Buffalo & Coop, 2020) demonstrated how background selection caused genome-wide positive temporal autocorrelation in allele frequencies (i.e. directional allele frequency change) as deleterious variants are selected against. Thus, conserved genes might show some evidence of directional allele frequency change. However, it is unknown how purifying and background selection impact locus-specific temporal change.

Here, we sequenced 130 herbarium specimens of *Arabidopsis thaliana* and combined these with new sequencing of recent field collected accessions and seedbank accessions, and published sequences of natural inbred lines and herbarium specimens (Alonso-Blanco et al., 2016; Durvasula et al., 2017). These genotypes greatly expand the geographic range of sequenced Arabidopsis genomes into regions of Africa and Asia, and were collected from nature across a period of nearly 200 years. These data were used for several goals. First, we sought to generate a more complete view of Arabidopsis genetic diversity and population structure.

Second, we sought to determine how population genetic composition has changed over time across the species range. Toward these goals we focused on the following questions:

1. What is the geography of population structure and relatedness across Africa and Eurasia? Are African populations diverse and genetically distinct from each other and from those in Eurasia (i.e. “relicts”)?
2. Has geographic population structure remained static? Or have new lineages spread across the Arabidopsis range over the last 150 years?
3. Is there genetic turnover through time, indicating isolation-by-time?
4. Are QTL for ecologically important traits enriched for temporal turnover in allele frequency, suggesting changing selection?
5. Do conserved genes show lower temporal turnover than non-conserved loci, suggesting continued conservation? Or do conserved genes show higher turnover due to purifying and background selection?

## Methods

### Sampling of herbarium specimens, wild individuals, and published data

We obtained genomes of 527 *Arabidopsis thaliana* samples for this study (Table S1). This includes data newly generated (n=225) from multiple sources: herbarium specimens with ancient DNA (aDNA) protocols in a clean lab (n=130), new collections from extant wild populations (n=91), and stock center accessions of (n=4) naturally inbred lines. We also used 302 published genomes from stock center lines (n=199), herbarium (n=38) and natural populations (n=65) (Alonso-Blanco et al., 2016; Durvasula et al., 2017; Lang et al., 2024). The published genomes from stock center lines were mostly collected in the last two decades (Figure 1). However, a few dozen were originally collected in western Europe in earlier decades. For temporal analyses, these accessions were included based on the year they were collected in the wild, following (Wilczek et al., 2014).

**Figure 1.**
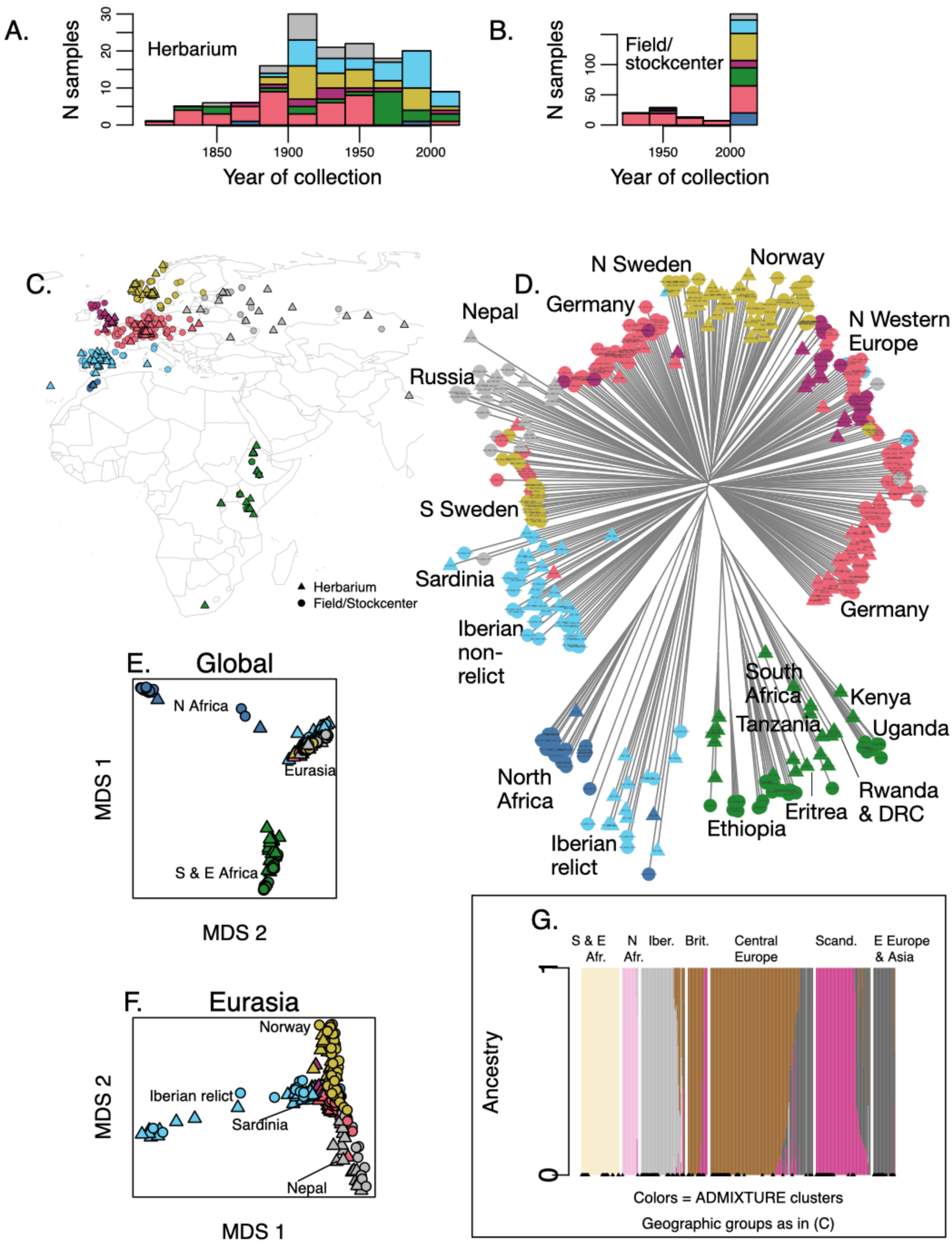
Sequenced accessions showing their distribution over time (A-B) and colored according to geography (C) or ADMIXTURE cluster (G) and with shapes indicating whether they were herbarium specimens (triangles) or sequenced from stock center or newly collected accessions (circles). Neighbor joining tree (D) includes sample codes with country abbreviation and year collected from the wild (Table S1). Two-dimensional MDS for the entire dataset (E) highlights the divergence between Africa and Eurasia and (F) for Eurasia highlights the distinct nature of the Iberian relicts. Specimens from countries of note are labeled on the tree. Cross validation error lowest for ADMIXTURE K=6 so these results are shown (G), with triangles at bottom indicating herbarium specimens. Shown are only accessions included in population genetic analyses (after filtering for balance between modern collections and herbarium).

Herbarium sampling was greatest for Germany, Spain, and Norway (n>30 accessions each), with additional sampling from Russia, Ethiopia, and Britain (n>10 accessions each). To uncover population genomic patterns in unstudied regions, we included samples from regions previously unrepresented in the literature: 5 countries (Kenya, Rwanda, Eritrea, the Democratic Republic of Congo, Nepal) and a large Mediterranean island (Sardinia). We also sequenced new populations from north of the Arctic circle, up to >68° latitude in Norway. The herbarium samples ranged in year of collection from 1817 (Germany), 1838 (Ethiopia), 1860 (Russia) to as recent as 2011 (Tanzania) and 2010 (Spain). To compare with herbarium specimens from the same region, we made new field collections of 91 individuals, 21 from Ethiopia (in 2017/2018), 9 from Uganda (in 2018), and 61 from Norway (in 2009/2010).

Publicly available data were obtained from 199 accessions of the 1001 Genomes project, selected based on geographic proximity to herbarium specimens (Alonso-Blanco et al., 2016) and from 70 African samples (65 individuals from wild populations and 5 herbarium specimens) of (Durvasula et al., 2017). In total, our samples encompass 32 countries in Eurasia and Africa, representing a large part of the native range of Arabidopsis (Figure 1) (Yim et al., 2023). We then down-sampled the modern collections from Morocco and Norway to balance with the sampling from herbarium specimens.

### DNA extraction, library preparation and sequencing: Newly generated data

130 newly obtained samples from herbaria were transported into a dedicated ancient DNA clean laboratory at Pennsylvania State University, USA. DNA was extracted and using specialized protocols detailed in Supplemental Material. After DNA extraction, we investigated the DNA fragmentation pattern for each sample and proceeded to perform library preparation based on the particular pattern of each sample. Using the following criterion, samples were categorized as high or low DNA fragmentation. Lowly fragmented samples showed DNA above the 500 bp ladder and were subjected to a short shearing step (30 seconds) using a M220 Focused Ultrasonicator (Covaris) prior to library preparation (detailed library preparation protocol in Supplemental Material), while highly fragmented samples were unsheared.

A total of 95 fresh samples were included in this study (61 field samples from Norway, 30 from Africa and 4 from the INRA stock center). Thirteen fresh leaf samples were collected from the field in East Africa while the remaining 17 were collected as seeds. Details for DNA extraction and library preparation are found in the Supplemental Material. Samples were sequenced on Illumina platforms to obtain PE 150bp reads.

### Data preprocessing and de novo SNP calling

Full details on filtering, mapping, and SNP calling are found in the Supplemental Materials. Herbarium samples were mostly newly generated data (130). To these we added five samples from the African genomes (Durvasula et al., 2017) and 33 German genomes (Lang et al., 2024) also collected from herbarium vouchers. Raw data for the African genomes was obtained from the European Nucleotide archive (ENA) of the European Molecular Biology Lab - European Bioinformatics Institute (EMBL - EBI) and German genomes from (Lang et al., 2024). Raw read sequence data were assessed with FastQC (http://www.bioinformatics.babraham.ac.uk/projects) to confirm that they met our quality standards. Genome sequencing coverage ranged from 81.41% to 98.91%. Mean depth of the covered portion of the genome showed a wide range, from 1 to 42.6 (median=8.4, Table S2). For all herbarium samples cytosine deamination profiles characteristic of ancient DNA were verified using mapDamage 2.0 (Jónsson et al., 2013) (Figure S1). SNP discovery was done using a pseudohaplotype approach (see Supplemental Materials for more detail) (Kistler et al., 2018).

### Statistical analysis

#### Population structure

We first analyzed global population structure by generating a set of high coverage and unlinked SNPs, i.e. those that remained after excluding those with >5% missing calls and filtering for linkage disequilibrium (PLINK *indep-pairwise* filtering SNPs with R^2^>0.1 in 50 SNP windows with 10 SNP steps between windows) (Chang et al., 2015). We calculated pairwise genetic distances between all samples (identity-by-state/Hamming distance, PLINK, flat missingness correction) to generate a neighbor joining tree and multidimensional scaling k=2 plot. We also calculated genetic distances and MDS in two dimensions separately after repeating the missingness and linkage filtering for two regional sets: 1) east and south African genotypes and 2) Eurasian genotypes.

To further characterize population structure, we implemented ADMIXTURE genetic clustering, across a range of cluster numbers (1 to 15) (Alexander et al., 2009). We applied ADMIXTURE to all global samples combined as well as a subset of south and east African samples, the region sampled well by our new data but previously the least studied.

#### Isolation by distance

We tested for geographic population structure and isolation by distance in spatially explicit analyses. To account for non-independence among pairwise distance measures involving the same individual, we fit mixed-effects models that include a random effect for each individual. We then compared them with null models based on AIC (ResistanceGA, (Peterman, 2018), these results were consistent when we excluded samples with >25% missing SNP calls).

To better understand the spatial scales of isolation by distance and its heterogeneity among the previously well studied Eurasian populations compared to the newly sampled regions of Africa, we calculated wavelet genetic dissimilarity (Lasky et al., 2024) which quantifies scale-specific genetic turnover among populations. That is, wavelet dissimilarity is a measure of genetic distance at a specific spatial scale (determined by the dilation of the wavelet function). Wavelet genetic dissimilarity increases at greater scales under patterns of isolation by distance, patterns that may differ between regions based on their level of gene flow and population age (Lasky et al., 2024). Within Eurasia and two African regions, we calculated the average wavelet dissimilarity among sampled locations at a range of scales from 1-500 km.

#### Isolation by time

We studied whether geographic population structure was stable over the period of the study (1817-2018). To test for major turnover in genetic composition, we focused on the well-sampled regions in Europe and asked whether different genetic clusters emerged within regions over time. We implemented ADMIXTURE on Eurasian genotypes and chose K=6 because it resolved many regional differences in genotype before a sharp increase in cross-validation error at K=7 (Figure S3). We then divided the well-sampled areas into five discrete regions: Iberia, Britain, Norway, Northern Germany, and Southern Germany, each dominated by a different genetic cluster. We then used non-parametric Spearman’s rank correlations to test whether the proportional assignment of genotypes for the most dominant ADMIXTURE cluster within each region changed over time.

To test for more continuous temporal genomic change, we fit mixed effects models as with geographic distance, but added a covariate for separation in time (year of collection) to explain pairwise genetic distances. As with the pure geographic models, these models include individual-specific random effects to account for individuals that may have particularly low or high genetic distance compared to all others. For three regions that were the best sampled (Germany, Norway, and Iberia), we also tested for turnover through time within the region while accounting for geographic distance.

#### Change in allele frequency at specific loci

We tested for evidence of directional allele frequency shifts in QTL underlying traits that are potentially important in environmental adaptation in Arabidopsis and show evidence of temporal phenotypic change. We previously documented change in Arabidopsis herbarium phenotypes, including many of the sequenced specimens here (DeLeo et al. 2020). In particular, we found an increase in the date of collection and photothermal units accumulated by date of collection, signifying phenological shifts where plants were collected later in growing seasons across eastern Europe and central Asia (DeLeo et al. 2020). We also found an increase in leaf C:N ratio over time in a region from the Mediterranean to central Europe (DeLeo et al. 2020). While DeLeo et al. (2020) found no change in ^13^C discrimination (Δ^13^C) over time, Lang et al. (2024) recently showed changes in the frequency of alleles associated with stomatal traits over time, thus we decided to include putative QTL for Δ^13^C. We used mixed-model GWAS results for flowering time from (Alonso-Blanco et al., 2016), for germination/dormancy from (Martínez-Berdeja et al., 2020), and for leaf C:N and δ^13^C from (Gamba et al., 2024). For flowering time (the trait measured on the most genotypes), we further implemented regional GWAS separately for Iberia, Germany, and Fennoscandia (Norway, Sweden, Finland) (Alonso-Blanco et al., 2016). We used univariate linear mixed models in GEMMA v.0.98.3 (Zhou & Stephens, 2012) excluding loci with MAF < 0.05, calculating Wald test P-values.

To test for changes in selection over time at putative QTL for these traits, we first characterized temporal directional allele frequency trends across the genome. We used several analyses with different strengths. We first used non-parametric Wilcoxon tests determining if alternate alleles had different median years of sample collection (function ‘wilcox.test’ in base R) (R Core Team, 2023). Additionally, we used multiple logistic regression to test for allele frequency change over years while controlling for linear effects of latitude and longitude (function ‘glm’ in base R). Finally, we also included a test that included kinship random effects, to identify loci with patterns deviating from the genomic background change through time (Akbari et al., 2024), while also accounting for spatial trends associated with latitude and longitude (using “mmer” function in the “sommer” package) (Covarrubias-Pazaran, 2016).

We tested for these allele frequency temporal dynamics across the native range (Eurasia and Africa), and also within well-sampled regions (Iberia, Germany, Norway), because different regions may undergo different changes in selection or phenotypic change (DeLeo et al. 2020). We tested whether top GWAS loci were enriched for temporal allele frequency shifts. We determined the strength of temporal change in SNPs (defined by the bottom 0.05, 0.1, 0.25 *p*-value quantiles for the temporal allele frequency change tests) among the GWAS top SNPs (25, 50, or 100 SNPs, after removing nearby SNPs within 25 kb). We compared this quantile to those generated from a null distribution generated from 1,000 circular permutations (around the genome) of SNPs’ trait temporal change p-values.

#### Purifying and background selection

We tested whether there was evidence of distinct evolutionary processes acting on genic DNA versus non-genic DNA, and we also tested conserved versus non-conserved genes. First, we compared the 0.01 p-value quantile of temporal allele frequency change for genic versus non-genic markers. To generate a null distribution for this comparison, we conducted 1,000 permutations of annotation of genic versus non-genic (circularly around the genome).

Next, we used a list of Arabidopsis genes from orthogroups conserved across eudicots identified by (Sun et al., 2018): 16,799 conserved versus 10,856 non-conserved genes. Conserved genes were defined based on orthogrouping of all genes from Arabidopsis and six other eudicot species (divergence >120 mya, *Ipomoea nil, Solanum tuberosum, Solanum lycopersicum, Capsicum annuum, Coffea canephora, Mimulus guttatus*) (Sun et al., 2018). Orthogroups that contained genes from at least six of these seven species were defined as conserved orthogroups. Arabidopsis genes within these conserved orthogroups were classified as conserved, while those not in conserved orthogroups were classified as non-conserved.

We first compared the 0.01 quantile of temporal allele frequency changes between all genic SNPs in conserved versus non-conserved genes, using permutations of SNP annotation. The site frequency spectrum likely differs between genic and intergenic SNPs, and between SNPs in conserved versus non-conserved genes, potentially influencing power to detect temporal change. Thus, we also tested these genomic contexts for enrichment of temporal allele frequency changes for “rare” (0.05≥MAF<0.1) and “common” (MAF>0.4) SNPs in separate analyses.

We also conducted gene-level tests. Because longer genes tend to show greater conservation (potentially stronger purifying selection) than shorter genes (Lipman et al., 2002), we divided genes into bins based on amino acid lengths. The effect of length on conservation was evident in the data from (Sun et al., 2018): non-conserved genes’ median length was 226 amino acids while the conserved median was 416 (Wilcoxon test, p<10^-16^). To obtain a test statistic for temporal change of each gene, we calculated the lower 0.1 tail (or 0.25, results were qualitatively the same) p-values for temporal change for SNPs in the coding region. We then permuted gene conservation status circularly around the genome 1000 times and developed a null distribution of median difference in gene p-values between conserved vs non-conserved genes, matching for gene length (in five bins: <200, 201-400, 401-600, 601-800, and 801-1000 amino acids long). We then compared the null permutations to the observed in a two-tailed test for each gene length bin. Because little is known about how background selection might influence variation in temporal allele frequency change across the genome, we conducted forward genetic simulations of purifying and background selection (see Supplemental Material for more detail).

#### Genomic prediction of trait change

As an alternate approach to testing for directional evolution of ecologically important traits, we also used whole-genome prediction. Whole genome prediction models can work well for polygenic traits and assume an infinitesimal model of trait variation (Hill & Kirkpatrick, 2010) and have previously been successfully employed to predict local adaptation in Arabidopsis (Gienapp et al., 2017). To create a genomic prediction model, we used a training dataset of 421 sequenced *A. thaliana* genotypes (Alonso-Blanco et al., 2016). Each genotype had flowering time data at constant temperatures of 10°C (FT10) and 16°C (FT16) (Alonso-Blanco et al., 2016). The kinship matrix was generated with PLINK v1.90b6.26 (Chang et al., 2015) using a minor allele frequency filter of 0.05. The ‘kin.blup’ function from the rrBLUP package (Endelman, 2011) was used to conduct genomic predictions for flowering time. To assess the accuracy, or extent of overfitting in our genomic prediction model, we cross-validated our data for FT10 and FT16 by conducting tenfold genomic predictions. We then used multiple linear regression models to test for changes in predicted flowering times across years within each of three regions (Iberia, Germany, and Norway), while accounting for elevation, latitude, and longitude.

## Results

### An expanded view of Arabidopsis population structure in its native range

We first analyzed global population structure using a set of high coverage, unlinked 201,299 SNPs. An MDS plot in two dimensions separates genotypes from North Africa, East and South Africa, and Eurasia (Figure 1C). ADMIXTURE cross-validation error was minimized at K=6. At K=6 the samples split into North African, South and East African, Iberian, British/Central European, Scandinavian, and Eastern European/Asian groups (Figure 1E, see other K values in Figure S2).

The Sardinian sample was from low elevation near the coast and was assigned to a mix of Eurasian non-relict and North African/Iberian relict ancestries. The Nepalese sample was estimated to have primarily Eurasian non-relict ancestry, but with small portions of ancestry from both African groups, suggesting potential ancestry of a relict group. The MDS in two dimensions of Eurasian populations showed the Nepalese sample is toward one extreme alongside Russian, Kazakh, and Ukrainian populations (Figure 1D).

African Arabidopsis populations are understudied but clearly genetically distinct (Figure 1B, 1C) (Durvasula et al., 2017). Because these African populations are under threat from future climate change (Yim et al., 2023), and because the bulk of previously unstudied countries contributed by our herbarium samples were from East Africa, we conducted a focused analysis on African populations. A neighbor joining tree showed major differentiation among mountains (Figure 2A). This differentiation is consistent with the overall rarity of Arabidopsis and low suitability in low elevations in East Africa (Yim et al., 2023), suggesting very little gene flow between mountains over long periods. However, the Tanzanian (mostly lower elevation mountains), Mt Suswa (Kenya), and South African populations were more similar to each other than were genotypes across other individual mountains, suggesting greater or more recent gene flow in these more southerly regions (Figure 2). There were signals of geographic barriers, with Ethiopian populations on either side of the Rift Valley being most different from each other (Figure 2).

**Figure 2.**
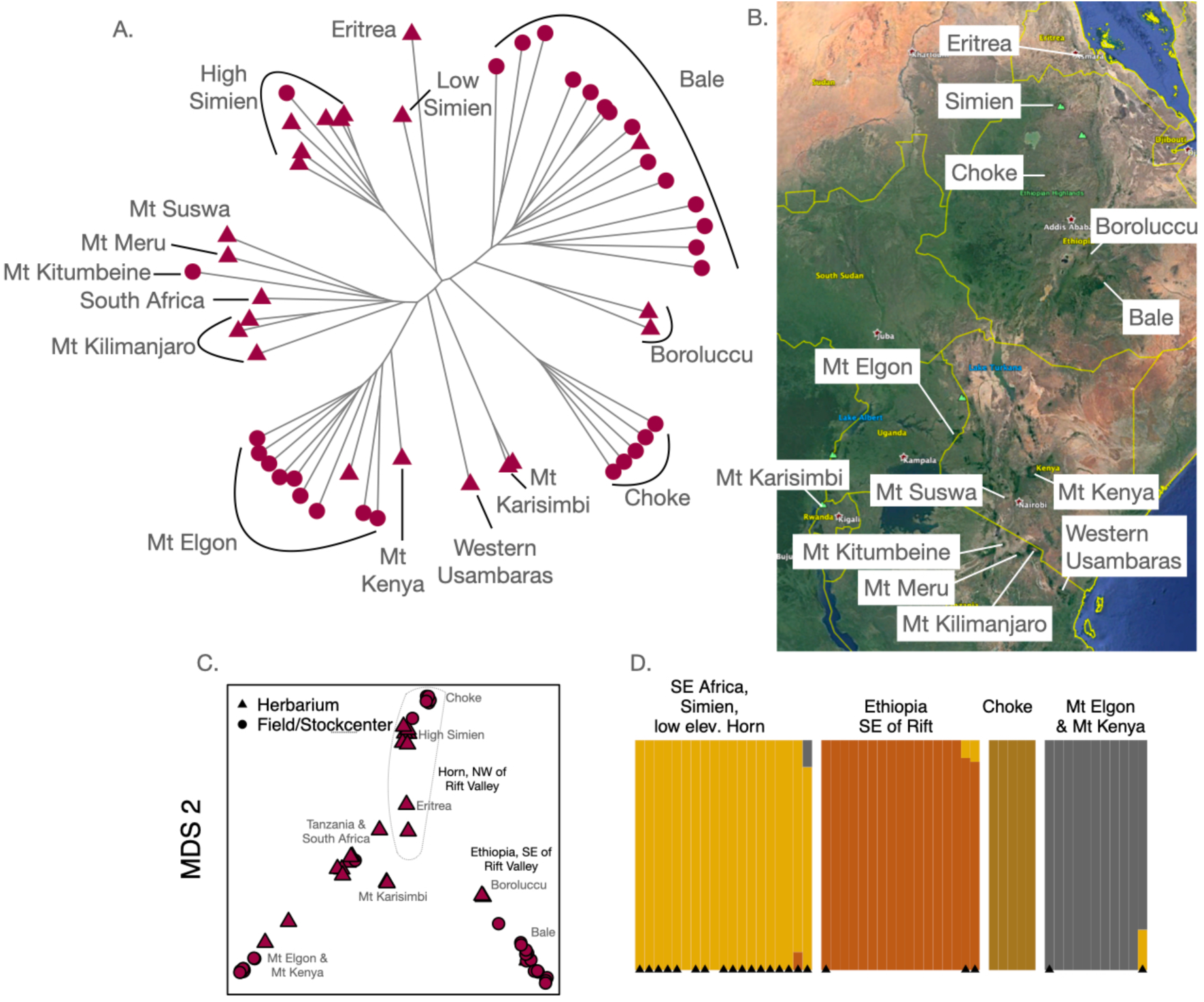
Accessions from East and South Africa, with shapes indicating whether they were herbarium specimens (triangles) or sequenced from fresh tissue from stock center or newly collected accessions (circles). Mountains of origin are labeled on the neighbor-joining tree (A) and map (B). (C) MDS in two dimensions with mountains labeled. (D) Cross validation error lowest for ADMIXTURE K=4 so these results are shown, with triangles at bottom indicating herbarium specimens.

An MDS plot of East/South African samples (n=52) divided Horn of Africa populations (Ethiopia & Eritrea) from those further south along the first dimension, while the second dimension separated populations on either side of the Rift Valley in Ethiopia from each other as well as those from further south (Figure 2C). ADMIXTURE K = 4 had the lowest cross validation error, splitting high elevation Ethiopian populations in Bale and Choke into two groups on either side of the rift valley, Mt Elgon & Mt Kenya shield volcanoes in Uganda & Kenya as another group. The Simien mountains, lower elevation samples in the Horn along with Mt Suswa (Kenya), all Tanzanian populations, Mt Karismbi (Democratic Republic of Congo & Rwanda) and South Africa were all in another group. The Rwandan & DRC samples are both from Mt Karisimbi, which straddles these nations’ borders and these two genotypes were highly similar.

### Patterns of population structure to assess the provenance of a historically important specimen

Genotypes from different elevations of the same mountain were typically more closely related than they were to genotypes from similar elevations on different mountains (Figure 2), as is true for most mountain ranges in Eurasia and Africa (Gamba et al., 2024). However, an exception was our oldest specimen in the region, from 1838, collected by Wilhelm G. Schimper at a location in the Simien Mountains. The location, according to Schimper, was “Demerki,” (sometimes “Demerkit”) a name not apparently in current use, or a corruption of another name. The “Demerki” sample falls in the neighbor-joining tree alongside a 1969 Eritrean sample from 1800 m asl (see “low elevation Simien” in Figure 2). ADMIXTURE on African accessions at K=5 (only slightly lower cross validation error than the K=4 shown in Figure 2) also assigns the ‘Demerki’ and Eritrean genotypes 30-40% ancestry from the group of genotypes from Tanzania, many at lower elevations (unlike the higher elevation Simien genotypes which show no such ancestry). Because of this similarity with lower elevation genotypes, we infer that “Demerki” was likely a site in Simien at lower elevation than 3500 m.

Based on the low genetic similarity, the demographic history of these Horn of Africa populations <2500 m asl (including populations known from herbarium specimens from Djibouti & Somalia) (Yim et al., 2023) appears distinct from the mostly higher elevation populations elsewhere in East Africa. Similarly, the 1953 sample from 1750 m asl in the Western Usambaras mountains in Tanzania was highly genetically differentiated from the other Tanzanian populations at higher elevations (Figure 2). Thus, in East African low mountain ranges near the Indian Ocean, Arabidopsis populations seem distinct from populations nearby at much higher elevations, possibly due to a distinct history, such as greater or more recent gene flow among populations, and adaptations to low elevations.

### Isolation by distance

We found significant isolation by distance globally including all accessions with coordinates (Figure 3, n = 404, linear mixed models with individual random effects, null AIC= -799279, distance AIC= -853089; general linear model of distance R^2^=0.41). For non-relict accessions in Eurasia, we also found significant isolation by distance (n=303, null AIC = -509158, distance AIC= -526778; general linear model of distance R^2^=0.36). Isolation by distance was also significant across East Africa (Figure 3, n=50, null AIC= -8998, distance AIC= -9765). Within the two most sampled East African mountains we also found significant isolation by distance: Mt Elgon, Uganda (max. distance = 18 km, n=10, null AIC= -432, distance AIC= -442) and the Bale Mts., Ethiopia (max. Distance = 43 km, n=14, null AIC= -821, distance AIC = -873).

**Figure 3.**
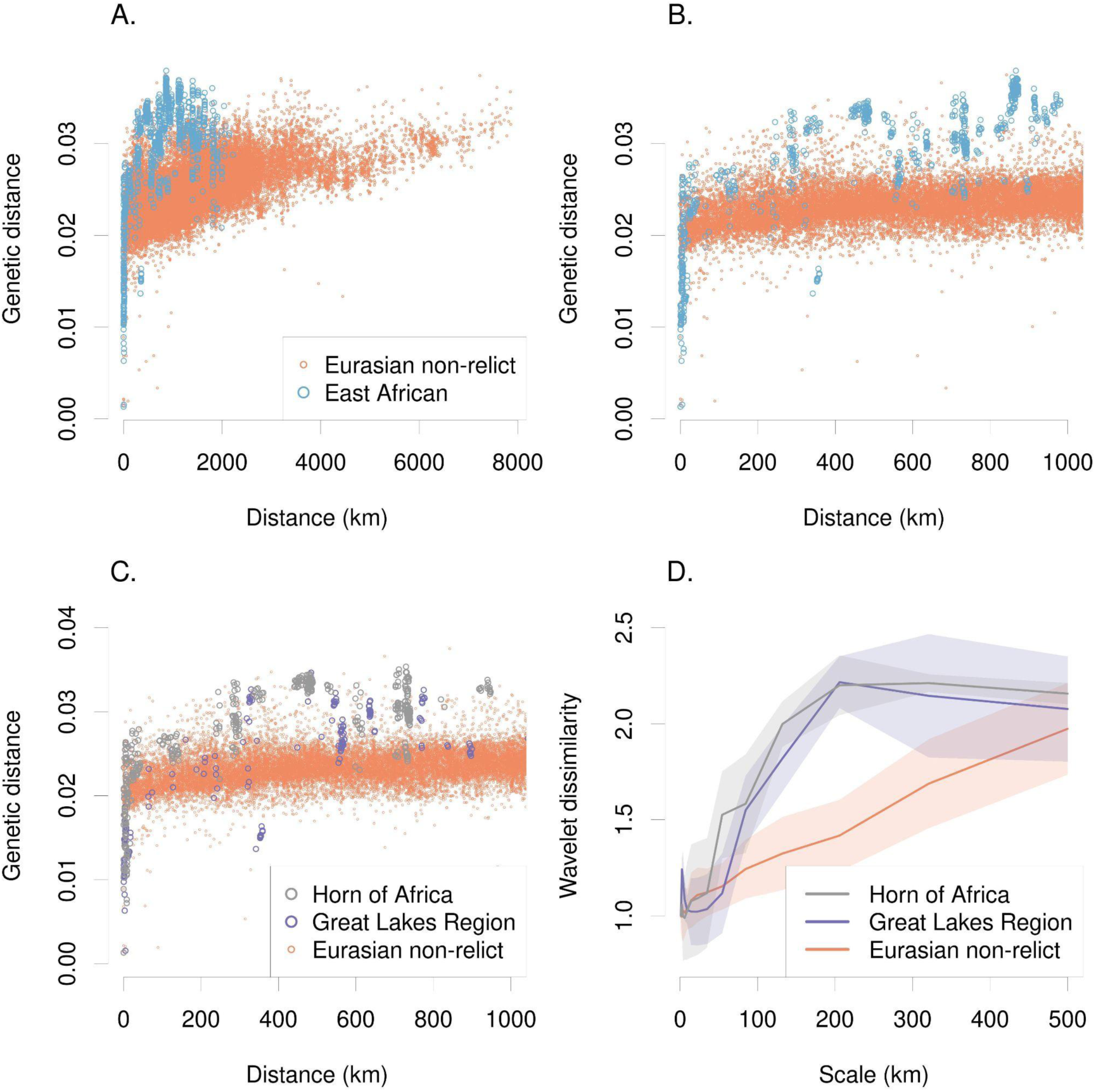
Isolation by distance across the range of Arabidopsis. (A) Pairwise genetic distances within regions, comparing Eurasian non-relicts to East African populations and (B) a truncated version focused on distances <1000 km. (C) Comparing pairwise genetic distance within two of the best sampled regions in East Africa with Eurasian non-relicts. (D) Scale-specific wavelet genetic dissimilarity (line = mean among sampled locations, ribbon = +-1 SD) showed the much greater genetic turnover at scales ∼100-500 km for both East African regions compared to Eurasian non-relicts.

Alonso-Blanco et al. (2016) found relict accessions showed a steeper increase in genetic distance over geographic distance, compared to non-relict accessions, suggesting longer histories of isolation for relicts. We compared East African samples to non-relict Eurasian samples and found a similar pattern: at distances > 250 km, East African samples were more genetically distinct than were non-relicts from Eurasia at the same distance (Figure 3). At a distance of ∼700 km we found the greatest genetic distance in East Africa; separating populations from the Great Lakes region from those farther north, in the Horn of Africa. These were more genetically distinct than were accessions from Nepal versus Portugal, >7800 km apart (the most geographically distant non-relict Eurasian samples). These results were also supported by wavelet dissimilarity analysis, which quantified scale-specific genetic turnover among populations. Horn populations showed the greatest turnover at ∼50-500 km scales, with Great Lakes populations just slightly lower (Figure 3). By contrast Eurasian non-relicts showed much lower genetic turnover up to ∼500 km scales.

### Genetic change through time and space

We saw no dramatic changes in ancestry over time. This can be observed in the neighbor-joining tree and ordinations (Figure 1) where herbarium specimens from a given region are most closely related to modern collections from the same region. Additionally, genetic clustering of Eurasian accessions by ADMIXTURE with K=6 showed little change in estimated ancestry assignments over the study period (Figure 4). None of the regionally dominant genetic clusters changed in the proportion of sample composition over time (Spearman’s rank correlation, year versus proportion of most regionally dominant cluster, all p>0.08; also true when using K=3, the K with lowest cross validation error in Eurasia). This indicates there were no major turnover events in population composition over the period of our study.

**Figure 4.**
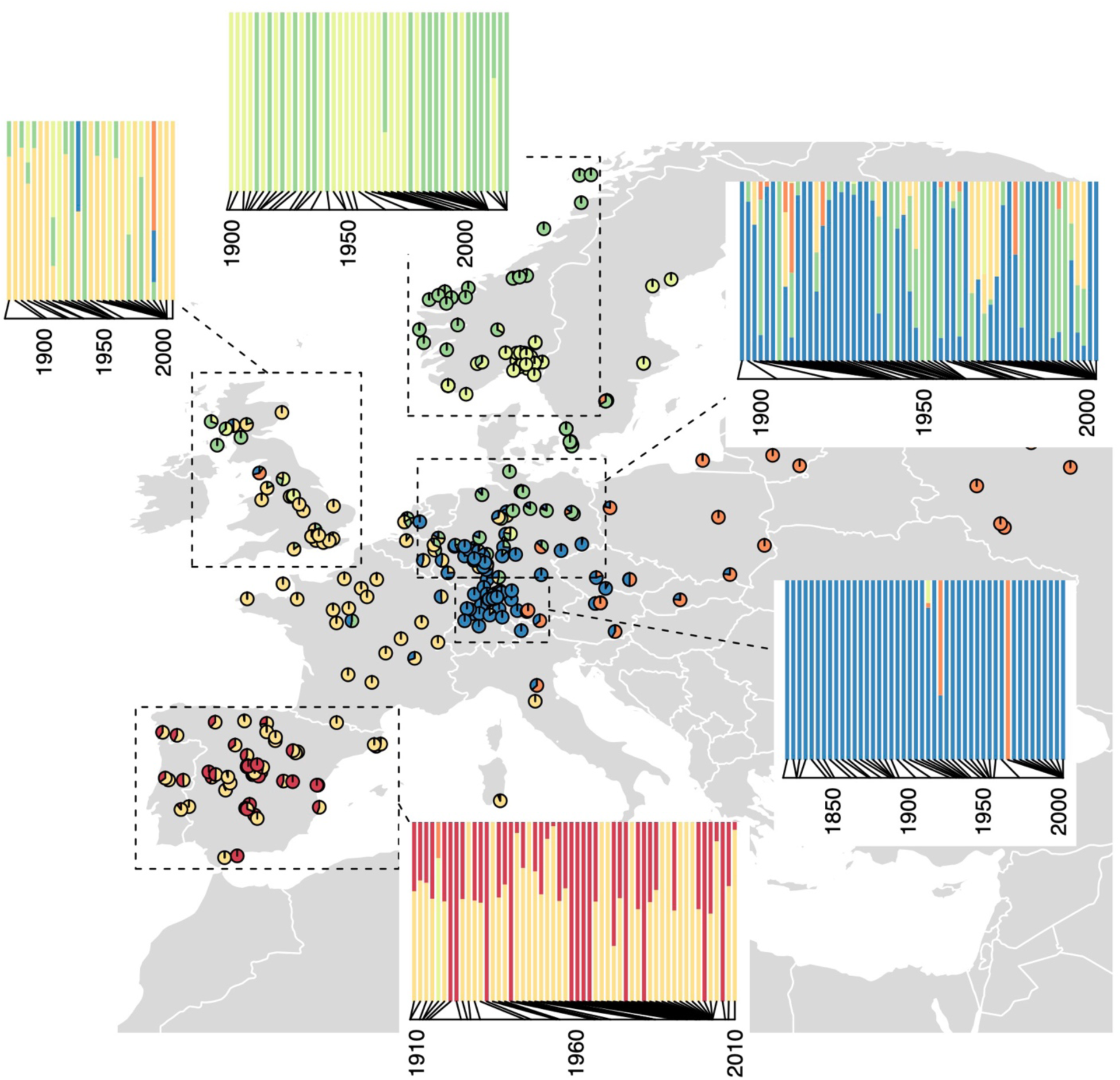
Genetic cluster membership over time (inset barplots) for regional subsets of genotypes (dashed boxes) with individual genotypes shown as pie charts indicated cluster assignment by ADMIXTURE with K=6. Each genetic cluster that was most common in a region showed no significant change in the proportion of assigned ancestry for local genotypes over time (Spearman’s rank correlation test, all p>0.08). Herbarium specimens and stock center genotypes are included here.

To examine changes in the distributions of deeper levels of genomic differentiation, we tested for changes in the distribution of relicts in Iberia, where they were at modest frequency (13%, Alonso-Blanco et al. 2016). Alonso-Blanco et al. (2016) noted that Iberian relict accessions in the 1001 Genomes panel tended to be found in landscapes with low anthropogenic disturbance and dry summers. We found in our Iberian herbarium sequences that this pattern was consistent across the 20th century (Figure 5B). Dividing Iberian samples into those from 1908-1968 and those from 1971-2010 (leaving equal sample size groups), and adding the 1001 Genomes genotypes (Alonso-Blanco et al., 2016), we found that relicts were in significantly drier summer climates but with no significant change in this pattern over time (precipitation in warmest quarter two-way ANOVA, relict p = 0.0358, time group p = 0.5111, relict X time group interaction p = 0.9458).

**Figure 5.**
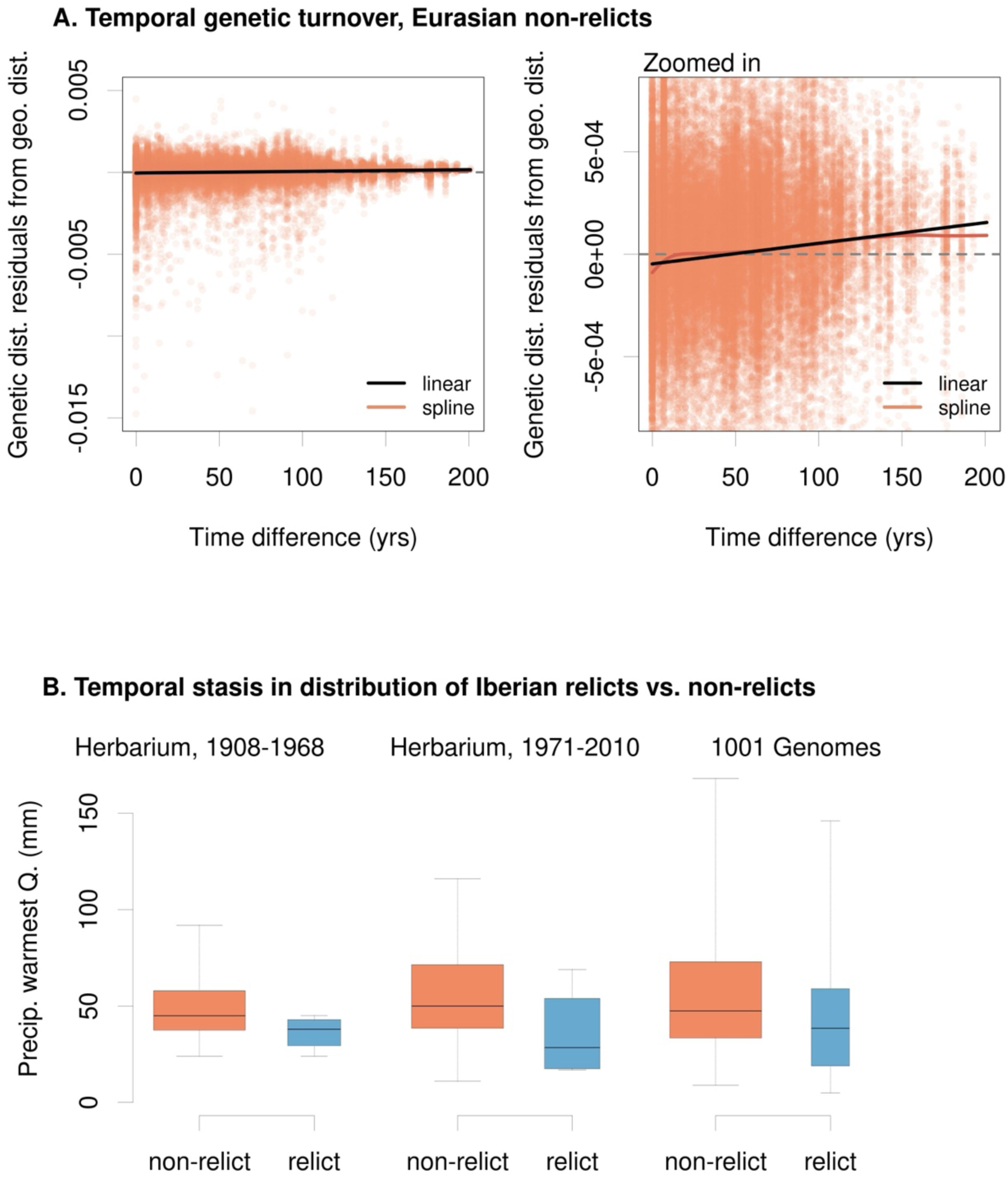
There is modest temporal genetic turnover (A) shown through residuals from a geographic mixed model vs. time difference, with a linear model (black) and spline (red) fitted, while (B) at coarser levels of relatedness the distribution of relicts in drier parts of Iberia remains consistent through time. Note the changing y-axis scales in (A) to show the statistically significant, albeit noisy, temporal turnover. (B) excludes Pyrenees because we did not have these populations sampled from herbaria.

Despite the stability in regionally dominant genetic clusters, we found evidence for modest continuous genetic turnover through time (Figure 5A). Among all regions, AIC favored a model of isolation by distance and isolation by time (n= 404, linear mixed model with distance only AIC = -853089, distance and time AIC = -853491; general linear model with distance only R^2^=0.41, distance + time R^2^ = 0.42). However, changes in genetic distance with time separation could partly result from changes in our relative sampling of different regions (e.g. 11% of herbarium samples before 1933, the median year, were relict lineages, vs. 32% 1933 and onward) or geographic heterogeneity in temporal change. To avoid this problem, we focused the analysis on some genetic clusters and geographic regions.

Among Eurasian non-relicts, a model with both time and geographic distance was again favored over a geography model only (n=319, linear mixed model with geographic distance only AIC = -577470, geographic and time distance AIC = -577688; general linear model with distance only R^2^=0.36, distance + time R^2^ = 0.38). To put the genetic turnover through time into a comparable context, we calculated the increase in genetic distance with 100 years separating samples in terms of the geographic distance expected to give the same genetic distance. While at first glance the slope of temporal genetic turnover in Figure 5 may seem slight, the estimated turnover across 100 years in Eurasian non-relicts was equivalent to that occurring over 185 km of geographic distance (for all accessions the figure was 299 km).

In Germany (n = 94), the model with isolation by distance and time was favored, such that with 100 years separating samples the genetic distance was expected to be of the same magnitude as the genetic distance between samples 70 km apart in geographic distance (mixed model geography AIC=-50286, geography + time AIC=-50314; general linear model with distance only R^2^=0.12, distance + time R^2^ = 0.13). For Norwegian (n=44) and non-relict Iberian samples (n=44) a model of isolation-by-distance (but not time) was favored by AIC (Norway: geography AIC=-9496, geography + time AIC=-9496; non-relict Iberia geography AIC=-9835, geography + time AIC=-9833).

### Temporal change in allele frequency of putative quantitative trait loci

In general, we found little consistent evidence that putative QTL for several ecologically important traits (GWAS SNPs) were enriched for directional allele frequency change. A list of top SNPs showing directional allele frequency change and nearby genes is included (Table S3 For Eurasian genotypes (n=339; 1,209,375 SNPs) several traits showed nominally significant (α=0.05) enrichments for some combinations of top GWAS SNPs and allele frequency change statistical model, but none of these were significant after controlling for FDR=0.05 (Table S4). The same was true in regional analyses using species-wide GWAS (none significant at FDR=0.05, Tables S5-7). However, because the genetic basis of variation may change among regions, we also tested SNPs from regional GWAS. We found Iberian flowering time (10°C) SNPs were strongly enriched in temporal allele frequency turnover (n=58 genotypes, top 25 SNPs, FDR=0.05, Table S8, Figure 6). We did not find any enrichment in Fennoscandia flowering time SNPs’ turnover in Norway (n=45 genotypes, Table S9). By contrast, German flowering time GWAS SNPs showed significantly reduced change over time, for both 10°C and 16°C (n=96 genotypes, Table S10, Figure 6).

**Figure 6.**
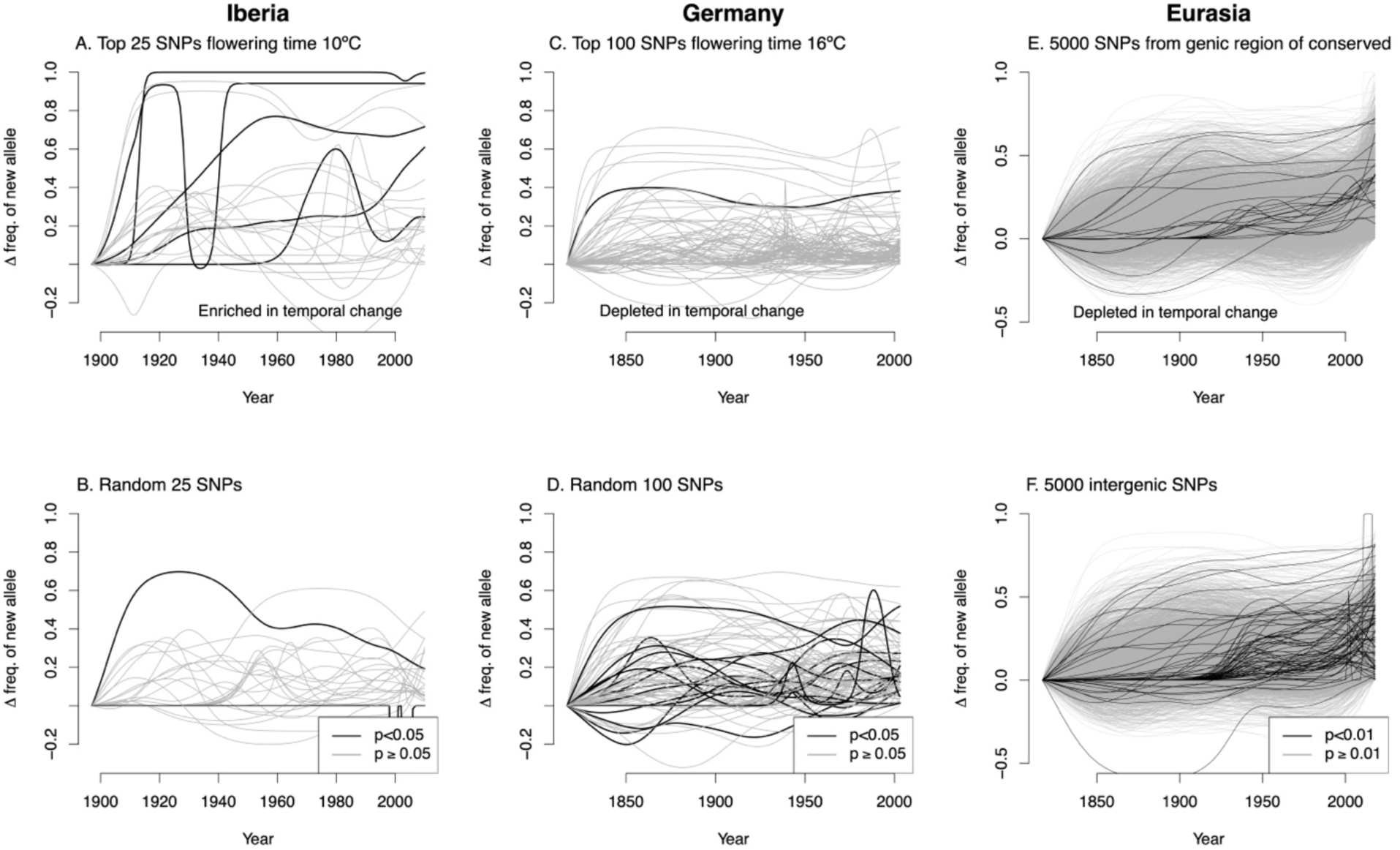
Allele frequency trajectories for different subsets of SNPs (lines) and regions, modeled using generalized additive models (GAMs) for visualization. Allele frequency trajectories are signed so that the y-axis indicates the increase in frequency of the allele more common in recent years (the ‘new allele’) compared to the beginning of the time series. SNPs with significant temporal allele frequency change in linear mixed models controlling for kinship and geography (p<0.05 for A-D, p<0.01 for E-F) are shown in black. The random SNPs and the genic/intergenic SNPs shown are single draws from the indicated categories.

### Lower directional allele frequency change in genes, especially conserved genes

Next, we tested enrichment of SNPs in genic regions for directional allele frequency change. We found across Eurasia, genic SNPs showed significantly less directional allele frequency change than intergenic SNPs genes (permutation test for 0.01 quantile of logistic and mixed model year effects, p=0.006 and p<0.002, respectively, example in Figure 6E-F). Additionally, genic SNPs of conserved genes showed significantly less directional allele frequency change than genic SNPs of non-conserved genes (permutation test for 0.01 quantile of logistic and mixed model year effects, p<0.002 for both). These patterns were generally consistent within regions (Table S11). These results were consistent when we restricted the analysis to only SNPs with MAF 5-10% and those with MAF > 40% (Table S12).

At the gene level, we also found that the conserved genes showed evidence for different types of selection depending on gene length (Figure S4). Eudicot conserved genes less than 200 AA sometimes had higher directional allele frequency change (p<0.05 for some models and regions, Table S13). By contrast, eudicot conserved genes from 200-400 AA were not significantly different from non-conserved genes. Furthermore, genes in amino acid length bins from 400-1,000 showed significantly less directional allele frequency change than null expectations (p<0.05 many models and regions, Table S13). We found these patterns were consistent within Germany and Iberia, but not statistically significant in Norway (Table S13).

The forward genetic simulations we conducted were variable in whether they showed genic regions under background selection having greater or less temporal allele frequency changes estimated with our approach (Figure S5). This variability is in contrast to the more consistent effect of background selection of increasing genome-wide temporal autocorrelation in allele frequency (Buffalo & Coop, 2020). In our simulations, genic SNPs (where deleterious mutations could occur; intergenic SNPs were all neutral) showed a significant depletion in temporal allele frequency changes in some simulations, while in other simulations they showed a significant enrichment in temporal allele frequency changes over intergenic SNPs.

### Temporal change in allele frequency: genomic prediction

Out-of-sample genomic predictions of flowering time were accurate with 10-fold cross validation (FT10 *r*=0.70, FT16 *r*=0.68). As a positive control, in genome-predicted flowering times we recovered a known cline: Iberia showed strong elevational clines (multiple linear regression; FT10 *t*=5.3, p<10^-6^; FT16 *t*=4.8, p<10^-5^) as observed from common garden experiments (Gamba et al., 2024; Montesinos-Navarro et al., 2011). When adding a year term to regressions, however, we only found a slightly significant advancement in predicted flowering time at 10°C over time in Germany (*t*=-2.5, p=0.0135) but otherwise we did not find any significant change in predicted phenology over time (Table S14). This general lack of clear evolution of flowering time is consistent with the collection date of herbarium specimens, which have not changed in western Europe, in contrast with eastern Europe to central Asia where dates were later in later years, regions not well sampled here (DeLeo et al., 2020).

## Discussion

To understand the importance of genetic variation in adaptation and phenotypic variation, it is important to characterize broader patterns of relatedness and genome-wide variation across a species. However, generating comprehensive population genomic surveys of widespread species is challenging due to the logistics of studies at such scales. For example, the Arabidopsis 1001 Genomes project made great progress by resequencing genotypes from across most of the European range (Alonso-Blanco et al., 2016), but greatly under-sampled regions of Asia and Africa (Durvasula et al., 2017; Hsu et al., 2019; Roy, 2018; Zou et al., 2017). Historically, research on Arabidopsis natural genetic variation has had a strong European bias (Hoffmann, 2002; Koornneef et al., 2004). However, African populations in particular show high genetic divergence from each other and from Eurasian populations (Durvasula et al., 2017; Gamba et al., 2024) and here we add substantially to the known diversity and its distribution in Africa. Additionally, high elevation African populations (samples in this study mostly from >4000 m asl) occupy outlier environments (Gamba et al., 2024), suggesting a great resource of undiscovered locally adapted genes and traits. Our use of museum sequencing allowed us to rapidly fill in important gaps in unstudied populations of Arabidopsis.

Temporal change in population size and allele frequency has been often inferred indirectly using current snapshots of population genetic composition (Nielsen et al., 2005; Schiffels & Durbin, 2014; Slatkin & Hudson, 1991). An exception is the rapidly growing field of ancient human genetics, which is starting to reveal the sometimes dramatic changes in population structure and ancestry over the last several thousand years (Allentoft et al., 2024; LaPolice et al., 2024; Olalde et al., 2018). However, there are few similar studies in other species outside of agricultural species (Kreiner et al., 2022; Smith et al., 2019; Swarts et al., 2017; Verdugo et al., 2019). By contrast, museums house millions of wild specimens covering centuries of sampling, providing a major resource to learn about evolution in response to contemporary environmental changes (Burbano & Gutaker, 2023; Lopez et al., 2020). Here, we used Arabidopsis specimen sequences to demonstrate temporal population genomic turnover, although we see little evidence of dramatic turnover in ancestry of the type seen in humans. Nevertheless, this turnover is heterogeneous across the genome, identifiably so, despite the limited outcrossing in this species. We showed how many genes conserved over many millions of years show distinct patterns of turnover during the study period (1817-2018).

### Global population structure

By sequencing museum specimens, we were able to fill in major gaps in global population structure especially in Africa, and important additions in Norway, Sardinia, and Nepal. We found that east and southern African Arabidopsis represent a group of populations highly distinct from those in North Africa and Eurasia, with major divergence between populations from different mountains across east Africa (Gamba et al., 2024). We showed that the level of genetic divergence between African populations increases with spatial distance at a rate much greater than that seen in Eurasia. This is consistent with the hypothesis that these populations have been spread across East Africa for a long period of time and represent a potential origin of the *Arabidopsis thaliana* species (Durvasula et al., 2017; Fulgione & Hancock, 2018).

The high divergence between mountains was especially true for high elevation populations in the Horn of Africa where the East African Rift Valley separates the two groups of greatest divergence, consistent with findings in *Trifolium* (Wondimu et al., 2014). Furthermore, the apparently high diversity of Arabidopsis from Bale (Figure 2) is consistent with these mountains also being a center of diversity for *Trifolium* and *Carduus* (Wondimu et al., 2014) perhaps because of the topography of Bale: a high elevation plateau with the widest range of Afroalpine habitat in the region. Within some of the relatively lower elevation populations in southern Kenya and northern Tanzania, we found less divergence suggesting more recent gene flow between populations. The separation of these genotypes from those nearby on Mt Kenya & Mt Elgon is consistent with findings in *Dendrosenecio* in this region (Tusiime et al., 2020). Our study helps set the stage for future investigation of adaptive and phenotypic variation in the diverse populations of East Africa.

The sample collected by W.G. Schimper from “Demerki” Ethiopia likely has a low elevation (<3000 m) provenance. Sources have disputed the location of the historically important “Demerki” specimens, as being Däräsge Maryam at 2990 m (Dorothea McEwan pers. comm.) or alternatively at 3500 m (Gillett, 1972), and many have georeferenced it at the peak of the massif (4400 m asl), though the specimen itself does not indicate elevation. The location is of interest given the extensive collections (>600 Schimper specimens labeled “Demerki” in GBIF), including many types, made by Schimper in a time period from which few other specimens survive (Gillett, 1972).

We also added a genome from a large Mediterranean island previously unstudied: Sardinia, which was a mix of Eurasian non-relict and Iberian/Moroccan relict-like ancestries. This relict ancestry is consistent with Sardinia being a refugium during glacial periods as recently indicated in our paleoclimatic projection of Arabidopsis distribution (Yim et al., 2023) and with previous findings that relict ancestry is greater in low elevations in the Mediterranean (Alonso-Blanco et al., 2016). Thus, it is likely that with greater sampling additional diversity of relict ancestry will be found across the Mediterranean. Previous studies had included very few sequences of Norwegian genotypes. Here we show that they are a group of relatively similar genotypes with similar to others from Northern Europe including Sweden, Britain, and Germany

### Temporal population turnover

One of the most powerful benefits of sequencing museum specimens is to be able to obtain longitudinal samples of populations through time at multiple locations and populations. Some recent examples of major turnover in ancestry include ancient humans in Europe (Allentoft et al., 2024; LaPolice et al., 2024; Olalde et al., 2018), *Centaurea* plants under expansion of a polyploid lineage (Rosche et al., 2024), and domesticated pigs following their introduction into Europe (Frantz et al., 2019). Notably, these examples either can travel large distances under locomotion (mammals) or wind and animal attachment (*Centaurea*). Indeed, using modeling based on temporal autocorrelation in allele frequency change (Simon & Coop, 2024) estimated that the great majority of human genetic turnover in European populations was attributable to gene flow, with some contribution from drift. In non-relict Eurasian Arabidopsis, we found more modest but detectable turnover through a period of 100 years equivalent to that estimated to occur between populations 185 km apart at the same point in time. This magnitude may seem high given Arabidopsis is primarily self-fertilizing and with little mechanism of seed dispersal. However, there is water dispersal by some genotypes using mucilage (Saez-Aguayo et al., 2014), and hitchhiking on humans, livestock, large mammals, and birds could play a role. Furthermore, individual Arabidopsis populations might show strong drift in landscapes where they colonize recently disturbed patches and potentially go extinct as succession proceeds (Baron et al., 2015; Lorts & Lasky, 2020). Further work is required to understand processes contributing to population genomic turnover. Additionally, a weakness of museum specimens is that they were not specifically sampled for this type of study, and so the heterogeneity in sampling through time may limit our ability to detect population genetic changes. Each region showed distinct spatiotemporal gaps in sampling (Figure 4), potentially limiting our ability to accurately characterize allele frequency and population structure dynamics. Furthermore, the need to remove only small parts of tissue from specimens and consequent relatively intensive lab work limits the ability to scale up sample sizes.

### Locus-specific turnover in allele frequency

Outside of experimental evolution with microbes, there have been relatively few previous time-series studies dissecting patterns of directional allele frequency change over many generations at specific loci across the genome (Akbari et al., 2024; Exposito-Alonso, Vasseur, et al., 2018; Lang et al., 2024; Le et al., 2022; Mathieson et al., 2015). Our sequence data allowed us to test for evidence of non-random patterns in allele frequency change. In general, we did not see much evidence for changes in the frequency of GWAS SNPs for traits, such as flowering time, d13C, and seed dormancy, known to be important to local environmental adaptation in Arabidopsis (and presumably under shifting selection with environmental change) (Dittberner et al., 2018; Gamba et al., 2024; Martínez-Berdeja et al., 2020; Stinchcombe et al., 2004).

There may be multiple reasons we did not see more such evidence of adaptive temporal allele frequency shifts. GWAS appear to work well in many Arabidopsis traits (Alonso-Blanco et al., 2016; Atwell et al., 2010; Baxter et al., 2010) so it seems likely the GWAS results used contained true QTL, although false positives and negatives may remain. Limited recombination in Arabidopsis may limit resolution to see temporal changes across the genome, but between many individual Arabidopsis populations there are many past recombination events that limit LD and provide resolution for GWAS. Furthermore, whole genome predictions on flowering time at two temperatures did not show genome-wide shifts in these traits. One plausible explanation may be that while traits of plants in nature have changed over time (DeLeo et al., 2020), the shifts in selection and genetic architecture of traits may be sufficiently heterogeneous among populations to limit our ability to see evidence of adaptation across regions and continents (Gamba et al., 2024; Lopez-Arboleda et al., 2021). Finally, the method of looking at enrichment in GWAS SNPs for temporal allele frequency shifts may be of limited power. Our strategy for testing for adaptation at these loci is similar to that of (Akbari et al., 2024), although we do not rely on determination of a significance threshold for calling individual loci as under selection. In Arabidopsis, (Lang et al., 2024) developed a polygenic score for individual stomatal genes to show shifts in allele frequency suggesting shifts to decreased stomatal density, a potential strategy to be implemented in the future for more traits.

We did identify the top flowering time GWAS SNPs in Iberia as enriched for shifts in allele frequency over time, potentially signifying changes in selection. Flowering time is a trait often under shifting selection among environments in many species (Ågren et al., 2017; Munguía-Rosas et al., 2011) and has been identified as a trait showing temporal change following environmental events (Franks et al., 2007). However, it is hard to identify the specific change in selection pressures potentially acting in this case. While climate has changed in Iberia over this period (Corte-Real et al., 1998; Esteban-Parra et al., 1998), land use may also influence selection on flowering time in Arabidopsis. And there has been dramatic land use change, such as the dramatic abandonment of farmland in the mountains of Spain over the 20^th^ century, >90% in some regions (Lasanta et al., 2017).

The loci showing the strongest allele frequency turnover identified several genes with known roles in environmental response (Table S3). For example, the second SNP with the most significant change in Eurasia was closest to plant U-box E3 ligase 12 (PUB12) which plays a role in immunity and abscisic acid signaling (Kong et al., 2015; Yamaguchi et al., 2017). The top SNP in Germany was *EXTRA LARGE G PROTEIN 3* (*XLG3*) which is involved in pathogen associated molecular pattern signaling (Y. Wang et al., 2023). The number 5 SNP in Norway is in a cluster of *SMALL AUXIN UP RNA* (*SAUR26/27/28*) genes that regulate temperature responsive growth affected by known *cis*-regulatory variants and with allele frequency clines along temperature gradients (Z. Wang et al., 2019). These loci merit further investigation to determine the effects of this allelic variation.

### Purifying and background selection and temporal shifts in allele frequency

We found that genic regions and conserved genes with longer amino acid sequences showed low temporal allele frequency change over time, potentially due to purifying selection and background selection on linked sites. The former pattern stands in contrast to the theoretical genome-wide results of (Buffalo & Coop, 2020) who found that background selection causes temporal autocorrelation in allele frequency change across the genome. Regardless of the genome-wide pattern, in Arabidopsis there are clearly differences among genomic contexts in their temporal dynamics. It is notable that genic SNPs are enriched in signals of local adaptation in Arabidopsis (Hancock et al., 2011; Lasky et al., 2012, 2018), opposite to what we found for temporal dynamics, perhaps because temporal allele frequency changes across the genome are dominated by background selection and not adaptation to changing environments.

One hypothesis to explain our finding is the following. Alleles at appreciable frequency for our analysis (i.e. MAF>5%) in conserved genes are neutral, background selection limits their diversity within populations, but many of these neutral (e.g. synonymous) variants are fixed between populations. Over time, background selection limits change in the locally fixed alleles at these loci but the allele frequency in our analysis (due to variation between populations) remains the same. Thus low temporal allele frequency turnover in conserved genes. In less conserved (shorter) genes, variants of appreciable frequency are neutral or slightly deleterious (leading to consistent decrease in frequency as predicted by Buffalo & Coop 2020) or under positive selection potentially due to environmental change (leading to consistent increase in frequency). Relatedly, (Cvijović et al., 2018) showed in an asexual model how background selection leads to distinct trajectories of neutral allele frequency depending on initial frequency, and so differences among loci in the site frequency spectrum can lead to distinct temporal trajectories. However, more theory is required to develop predictions for differences in allele frequency trajectories across genomes subject to spatially variable purifying, background, and positive selection. The fact that conserved Arabidopsis genes were depleted in temporal allele frequency changes suggests Arabidopsis demographic and genome parameters may be in a space where background selection reduces temporal allele frequency change. Other species with different demography and genomes may show distinct patterns.

### Conclusions

Natural history collections contain a vast wealth of samples that can generate insights into how organisms and populations are changing over time. However, in recent years their survival has been threatened as institutions cut support. Our results highlight the continued vitality of these collections and potential benefits to directly revealing evolutionary change through time.

Arabidopsis is changing as likely all species, and future work will dissect ecological and evolutionary mechanisms driving this change.

## Acknowledgements

This research was made possible by assistance from several curators and museums, specifically Real Jardín Botánico, Kew Gardens, Komarov Botanical Garden, Oslo Museum of Natural History, Stuttgart State Museum of Natural History, and Herbarium Tubingense. Plant material was exported from Uganda with permission of The Ministry of Agriculture, Animal Industry and Fisheries’ Plant Quarantine and Inspection Services, permit UQIS 4414/93/PC (E). Material was exported from Ethiopia with permission of the Ethiopian Biodiversity Institute, Ref. no. EBE71/7065/2018. Material was imported to the USA under USDA APHIS permits P37-17-01651 and P37-18-00230. Sequencing of Norwegian accessions was supported by NSF DEB-1743273 to CGO and JKM. We thank D. Weigel and the Max Planck Society for funding the sequencing of German herbarium specimens. LL is partially supported by California State University, San Bernardino, and by National Science Foundation award BIO-BRC 2217793. Funding was provided by National Institutes of Health grant R35GM138300 to JRL.

## Author contributions

The study was conceived and designed by LL and JRL. LL, PL, AH, MY, PW, JK, CB, JRL collected data. LL, PL, SL, YX, EC, DG, and JRL analyzed data. LL and JRL led writing. All authors contributed to writing.

## Data accessibility

### Genetic data

Individual genotype data are available on DataDryad (DOI: 10.5061/dryad.31zcrjdz1). Raw reads are available at NCBI SRA as project number PRJNA1269104.

### Sample metadata

Metadata can be found in supplementary tables of this manuscript.

### Code

Scripts are available on GitHub (https://github.com/jesserlasky/Arabidopsis_herbarium) and on DataDryad (DOI: 10.5061/dryad.31zcrjdz1 ).

## Supplemental Material

### DNA extraction, library preparation and sequencing: Newly generated data

#### Herbarium samples

One hundred and thirty newly obtained samples from herbaria were transported into a dedicated ancient DNA laboratory at Pennsylvania State University, USA. To avoid introducing contamination, the plastic bags containing the samples were cleaned with a 2% bleach solution. The surface of each sample was gently cleaned of potential soil and dust particles with a thin brush. The brush was cleaned in between samples by submerging the tip in a 2% bleach solution and then thoroughly rinsing it with milli-Q water (Millipore Corporation). DNA extraction was done using one medium-size leaf of *Arabidopsis thaliana*. Prior to extraction, tissue was homogenized using a TissueLyser (Qiagen). Each sample followed two cycles of homogenization, the first cycle was done for 30 seconds and the second was 15 seconds to ensure all tissue had been pulverized. Then, the DNA was extracted using the PTB protocol recommended for ancient samples with minor modifications (Wales & Kistler, 2019). Modifications were made to the length of the digestion (24 hours), to the final amount of Buffer AW1 (5x) and the amount of the elution buffer (25 µl). After extraction, DNA concentration was determined in a Qubit fluorometer using the Qubit dsDNA high-sensitivity (HS) Assay Kit (ThermoFisher Scientific). In addition, 5 µl of DNA from each sample were run in a 1% agarose gel to visually inspect the degree of DNA fragmentation.

After DNA extraction, we proceeded to library preparation. DNA obtained from herbarium samples is expected to be highly fragmented. Thus, no shearing step is done before library preparation. However, since our study includes both samples from recent years (∼ 10 years ago) and samples collected one or two centuries ago, we performed library preparation based on the particular fragment-size of each sample. Samples that showed DNA above the 500 bp ladder band were subjected to a short shearing step (30 seconds) using a M220 Focused Ultrasonicator (Covaris). Samples lacking DNA above the 500 bp ladder band were processed for library preparation as recommended for ancient samples (no shearing step). Herbarium-preserved samples are known to show fragmentation and damage rapidly after preservation (Staats et al., 2011). Thus, to capture short fragments and maximize complexity despite low input concentration we followed a library preparation method that relies on blunt-end ligation of custom adapters (Meyer & Kircher, 2010). This method is known to be more efficient and less-biased for chemically damaged and fragmented DNA than the AT-overhang adapter ligation commonly used in commercially available library preparation kits (Seguin-Orlando et al., 2013). One modification was implemented for the non-sheared samples; cleaning steps during the library preparation process were done using first the QIAquickNucleotide Removal Kit (Qiagen) and then the MinElute PCR Purification Kit (Qiagen) instead of AMPure XP/SPRIselect magnetic beads (Beckman Coulter). Final DNA content in the libraries was quantified using a Qubit fluorometer with the Qubit dsDNA high-sensitivity (HS) Assay Kit (ThermoFisher Scientific). Since samples had individual identifying barcodes, a pooled equimolar aliquot was prepared for sequencing. Each sequencing run contained approximately 40 randomly selected samples, including extraction and library preparation blanks to ensure that no contamination had happened during any step. The 42 samples used for the pilot study were sequenced PE at 100bp in a HiSeq 2500 platform. The remaining samples were sequenced at 75bp PE in a NextSeq 550 Illumina sequencer.

#### Recent field collections

A total of 95 fresh samples were included in this study (61 field samples from Norway (McKay et al., 2025), 30 from East Africa and 4 from the INRA stock center). Thirteen fresh leaf samples were collected from the field in East Africa while the remaining 17 were collected as seeds. Leaf material was immediately desiccated in silica gel within sealed plastic bags. The samples were transported to the laboratory and stored at 4°C until DNA extraction. Seed samples from East Africa, together with seeds from the 4 INRA stock center lines, were cultivated in growth chambers, and leaf tissue was dried in silica gel. Prior to DNA extraction all samples were flash-frozen in liquid Nitrogen and then homogenized as done with the herbarium samples. Next, DNA was extracted using the DNeasy Plant Mini Kit (Qiagen) following the manufacturer’s recommendations. The extracted DNA concentrations were determined in a Qubit fluorometer using the Qubit dsDNA high-sensitivity (HS) Assay Kit (ThermoFisher Scientific). DNA from the field-collected leaf samples were sent to BGI (China) for library preparation and sequencing. These samples were sequenced with paired end (PE) 150bp reads on a HiSeq X Ten Illumina machine. The remaining samples were processed by Genomics and Bioinformatics Service at Texas A&M AgriLife, where library preparation was conducted and sequencing was done on a NovaSeq 6000 Illumina platform with PE 150bp reads. Finally, the Norwegian field samples were collected as seeds and cultivated in a Colorado State University greenhouse. A single plant was sampled from each family once plants reached the vegetative state, and leaf tissue was from that plant was used for subsequent DNA extraction using the Qiagen DNeasy Plant Mini Kit (Valencia CA, USA). Extracted DNA was then quantified using a Qubit Fluorometer (ThermoFisher Scientific). Whole genome sequencing (WGS) libraries of the extracted DNA were prepared at the University of Colorado Boulder sequencing core. These WGS libraries were then paired-end (2 x 150 bp) whole genome sequenced at the University of Colorado Anschutz Medical Campus using an Illumina HiSeq.

### Data preprocessing and de novo SNP calling

#### Herbarium samples preprocessing

Herbarium samples were mostly newly generated data (130). To these we added five samples from the African genomes (Durvasula et al., 2017) and 33 German genomes (Lang et al., 2024) also collected from herbarium vouchers. Raw data for the African genomes was obtained from the European Nucleotide archive (ENA) of the European Molecular Biology Lab - European Bioinformatics Institute (EMBL - EBI) and German genomes from (Lang et al., 2024). Raw read sequence data were assessed with FastQC (http://www.bioinformatics.babraham.ac.uk/projects) to confirm that they met our quality standards. For all herbarium samples cytosine deamination profiles characteristic of ancient DNA were verified using mapDamage 2.0 (Jónsson et al., 2013) (Figure S1). Samples processed using the traditional aDNA library preparation and samples sheared before library preparation were checked with MapDamage 2.0 separately to ensure that all meet the ancient DNA criteria. Reads were trimmed using Leehom with the -ancientdna flag (Renaud et al., 2014). Merged and unmerged reads were then concatenated to maximize genome coverage. Trimmed reads were mapped to the *Arabidopsis thaliana* nuclear genome (TAIR v. 10) with the BWA v.0.7.17 -*aln* function since this option has a higher mapping rate for short fragments compared with alternatives (Li, 2013; Li & Durbin, 2009). After the reads from each sample were mapped to the nuclear genome, SAM_TOOLS_ v.1.5 was used to transform the mapped SAM files to BAM, sort the BAM files, remove duplicates, and filter the reads (minimum mapping quality of 30 and 30 base pair minimum length) (Li et al., 2009). Finally, SAM_TOOLS_ *flagstat* was used to count the number of alignments for every sample mapped to the *A. thaliana* nuclear genome (Li et al., 2009).

#### Fresh samples and publicly available data preprocessing

Raw data from 199 samples (Alonso-Blanco et al., 2016) were downloaded from the Sequence Read Archive (SRA) of the National Center for Biotechnology Information (NCBI). Additionally, the 65 African samples (all Moroccan except one from Tanzania) were obtained from the European Nucleotide archive (ENA) of the European Molecular Biology Lab - European Bioinformatics Institute (EMBL - EBI). For all the publicly available samples and the ones generated from fresh tissue, raw read sequence data were assessed with FastQC (http://www.bioinformatics.babraham.ac.uk/projects) to confirm that they met our quality standards. Raw paired-end reads were trimmed and filtered using Trimmomatic-0.38 to remove adapter sequences, short reads, and low-quality reads from raw sequence data (Bolger et al., 2014). To remove adapter sequences, we used seed matches with a maximum of three mismatches and an initial length of 10. These seeds were extended and clipped when paired-end reads had a score of 30 or lower. Additionally, leading and trailing bases of low quality (Phred quality score <20) or N were removed. Likewise, reads where the quality of a 4-base-wide sliding window dropped below 15 at the 3’ end were removed. Finally, any remaining reads that were shorter than 75 bases long were eliminated. Trimmed reads were then mapped to the *Arabidopsis thaliana* nuclear genome (TAIR v. 10) with BWA v.0.7.17 -*mem* flag (Li, 2013; Li & Durbin, 2009). After the reads from each sample were mapped to the nuclear genome, SAM_TOOLS_ v.1.5 was used to convert the mapped SAM files to BAM, sort the BAM files, remove duplicates, and filter the reads (minimum mapping quality of 30 and 30 base pair minimum length) (Li et al., 2009). Finally, SAM_TOOLS_ -*flagstat* was used to count the number of alignments for every sample mapper to the *A. thaliana* nuclear genome (Li et al., 2009).

de novo *SNP calling -* SNP discovery was done using a pseudohaplotype approach (Kistler et al., 2018). A combination of SAM_TOOLS_ *mpileup* and VarScan v2.3.9 *mpileup2snp* was used to discover all candidate variant positions in all samples with minimum mapping quality of 20, a minimum read depth of 2 and at least 2 supporting reads to call a variant (Koboldt et al., 2009; Li et al., 2009). To exclude paralogs and repetitive regions, variable sites in the top 10% of coverage were filtered out and the remaining sites were summarized in a bed file. The bed file was used to create individual pseudohaplotype files for each sample of variable sites with a minimum coverage of 2 and a maximum of 70 (these coverage thresholds were selected based on the specific coverage data from all samples included in this study). After filtering sites by coverage, individual files were combined into plink tped and tfam files and culled to biallelic sites. Finally, PLINK v1.90 was used create the final vcf file containing the selected variable sites (Purcell et al., 2007).

#### Simulating purifying and background selection

We used the SLiM simulation software to simulate a Wright-Fisher model with semi-realistic genome structure (genes with introns and exons, and intergenic space) and a combination of neutral and deleterious mutations, with deleterious mutations only occurring in genic sequences (Haller & Messer, 2019). We simulated a stepping-stone model of 10 populations with a migration rate of 0.005 and used a selfing rate of 95%. We ran models for 100k generations, and tested simulations with different combinations of genome size, deleterious mutation distribution of fitness effects, and recombination rates. We then tested each SNP of at least 5% MAF in a sample of 200 individuals across the last 100 generation in regression models with time (as in the Arabidopsis data). We then tested for enrichment of the temporal allele frequency change statistics for mutations in different genomic contexts (genic vs intergenic).

---

**Figure S1:**
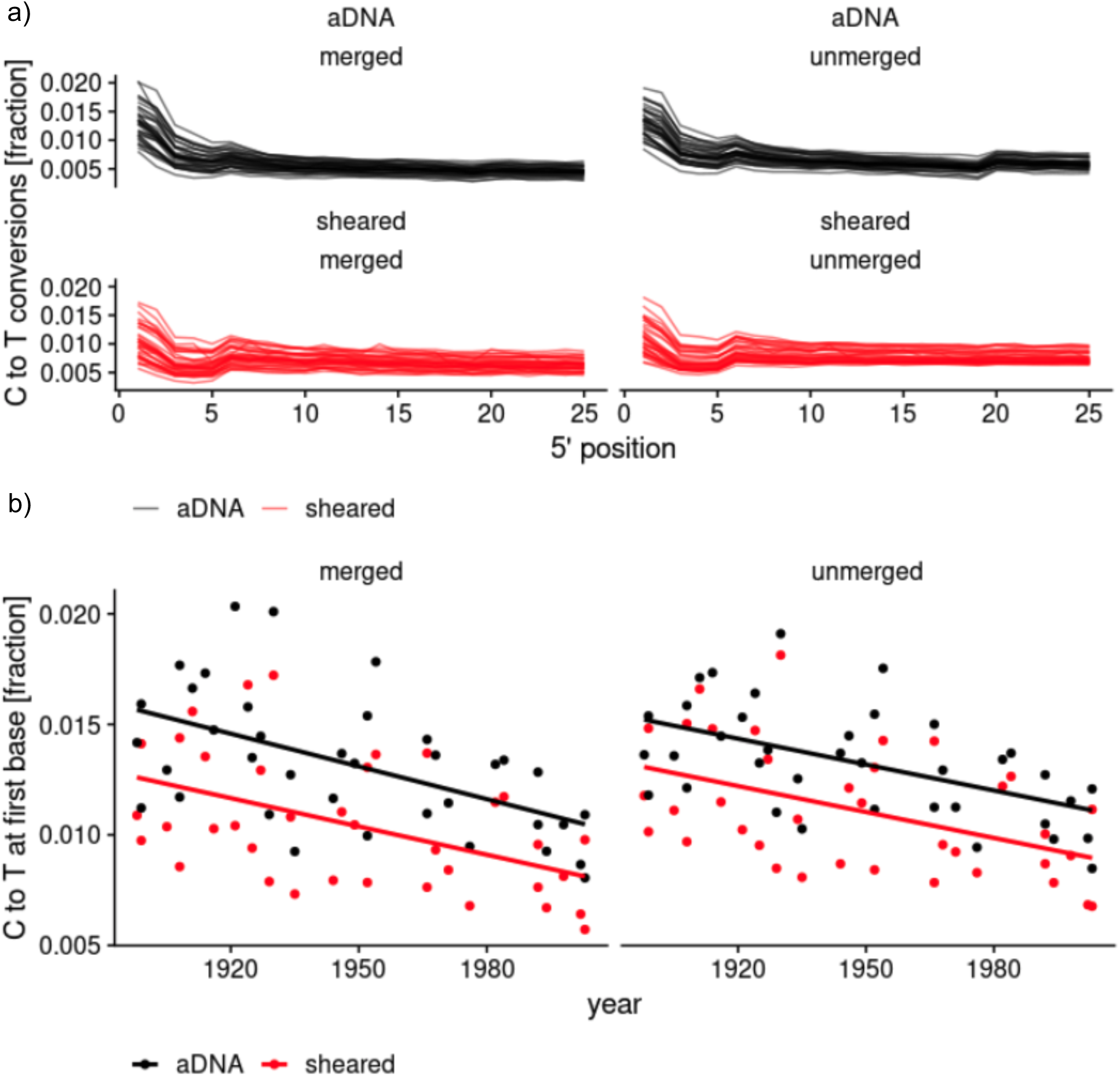
Historical sample authentication. Merged (on the left) and unmerged reads (on the right) were run separately. Black indicates samples that were not sheared (aDNA) and red indicates samples that were shared (sheared) before library preparation due to differences among samples in the existing degree of fragmentation. A) Fraction of C-to-T converted base-pairs along sequencing reads and the B) correlation of C-to-T conversion at the first base of a read respective to sample’s age. These patterns show the characteristic damage done to DNA fragments accumulated over time.

**Figure S2.**
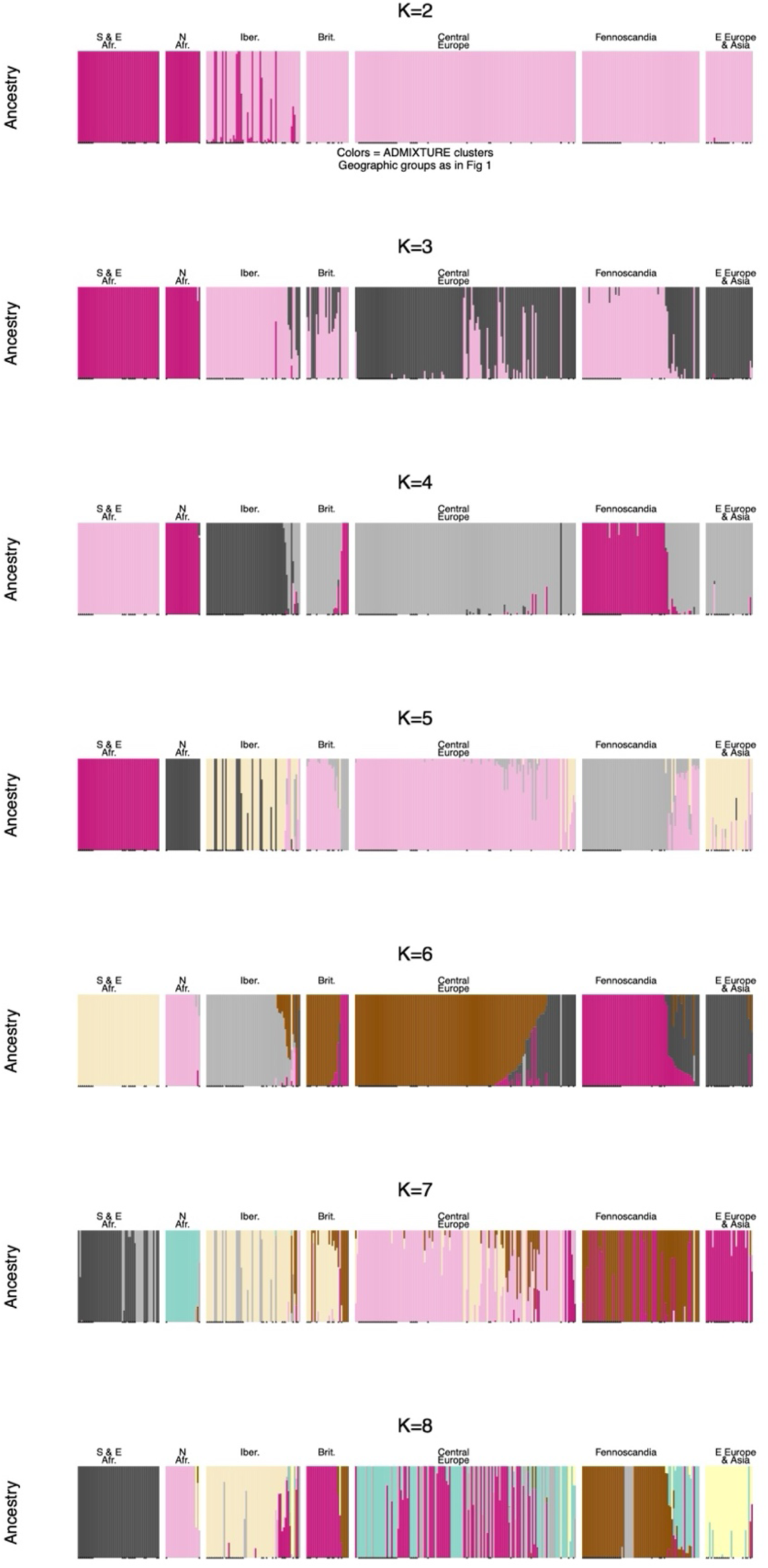
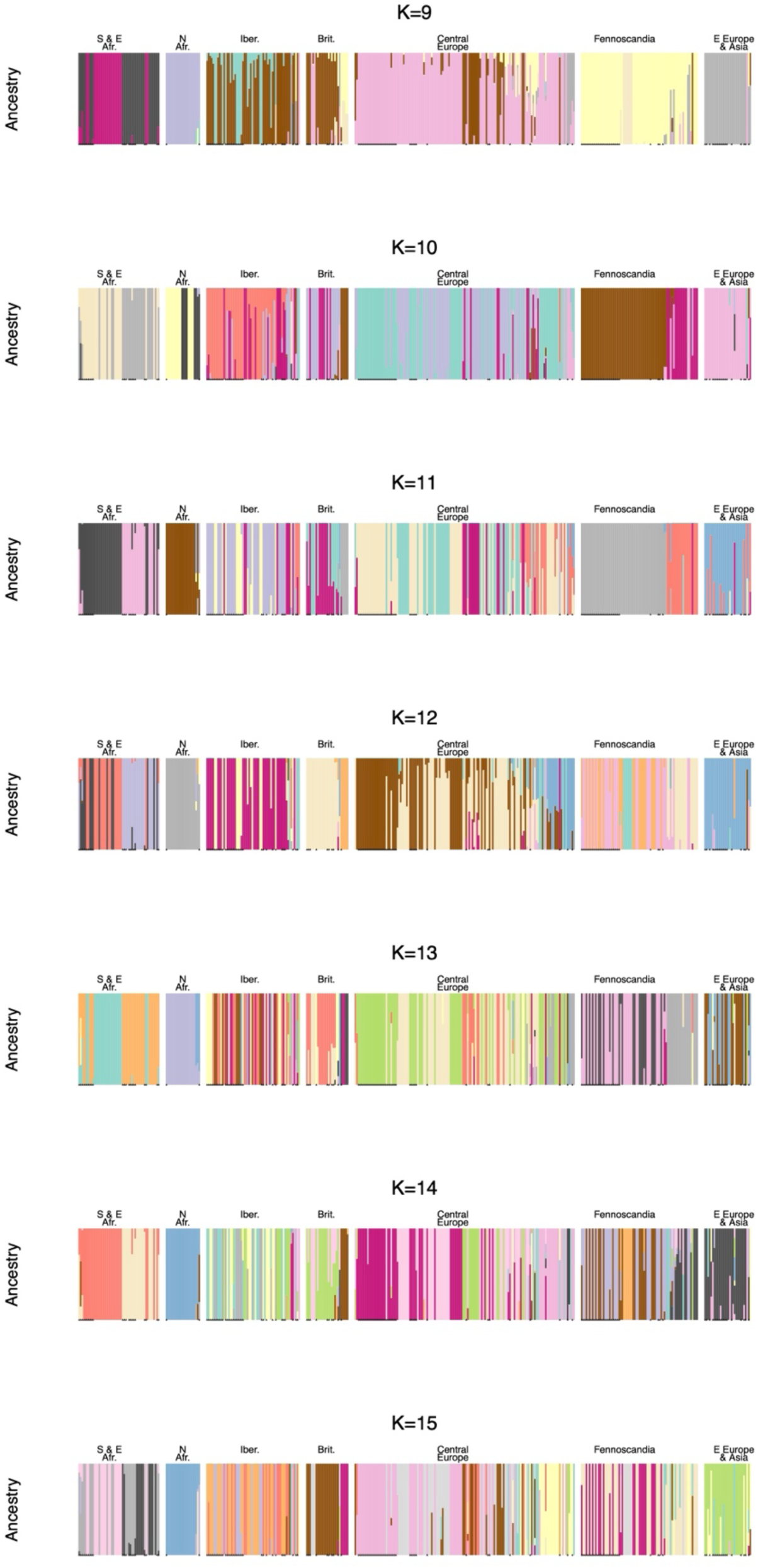
ADMIXTURE results for K=2-15, with samples grouped along the x-axis by geographic region as in Figure 1. Colors indicate assignment to ancestral clusters.

**Figure S3.**
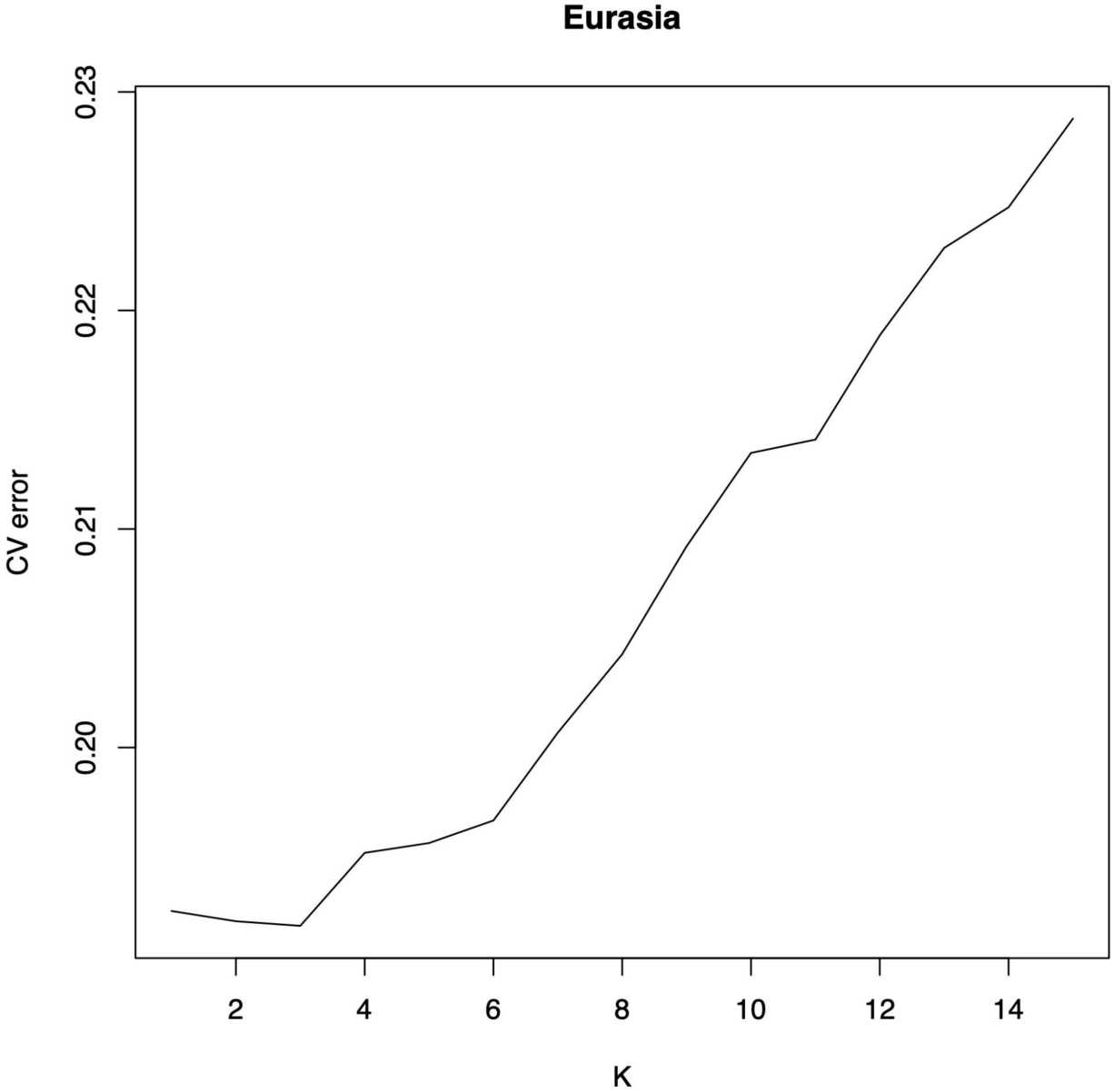
Results of ADMIXTURE clustering with different K values **across Eurasia**. This clustering was primarily for visualization purposes and determining turnover in genotypes within regions in Figure 4. For this reason, we used K=6 in analyses, though we also conducted a version of analysis with K=3. This K resolved different regional genotypic clusters but came before a sharp increase in cross validation error at K=7.

**Figure S4.**
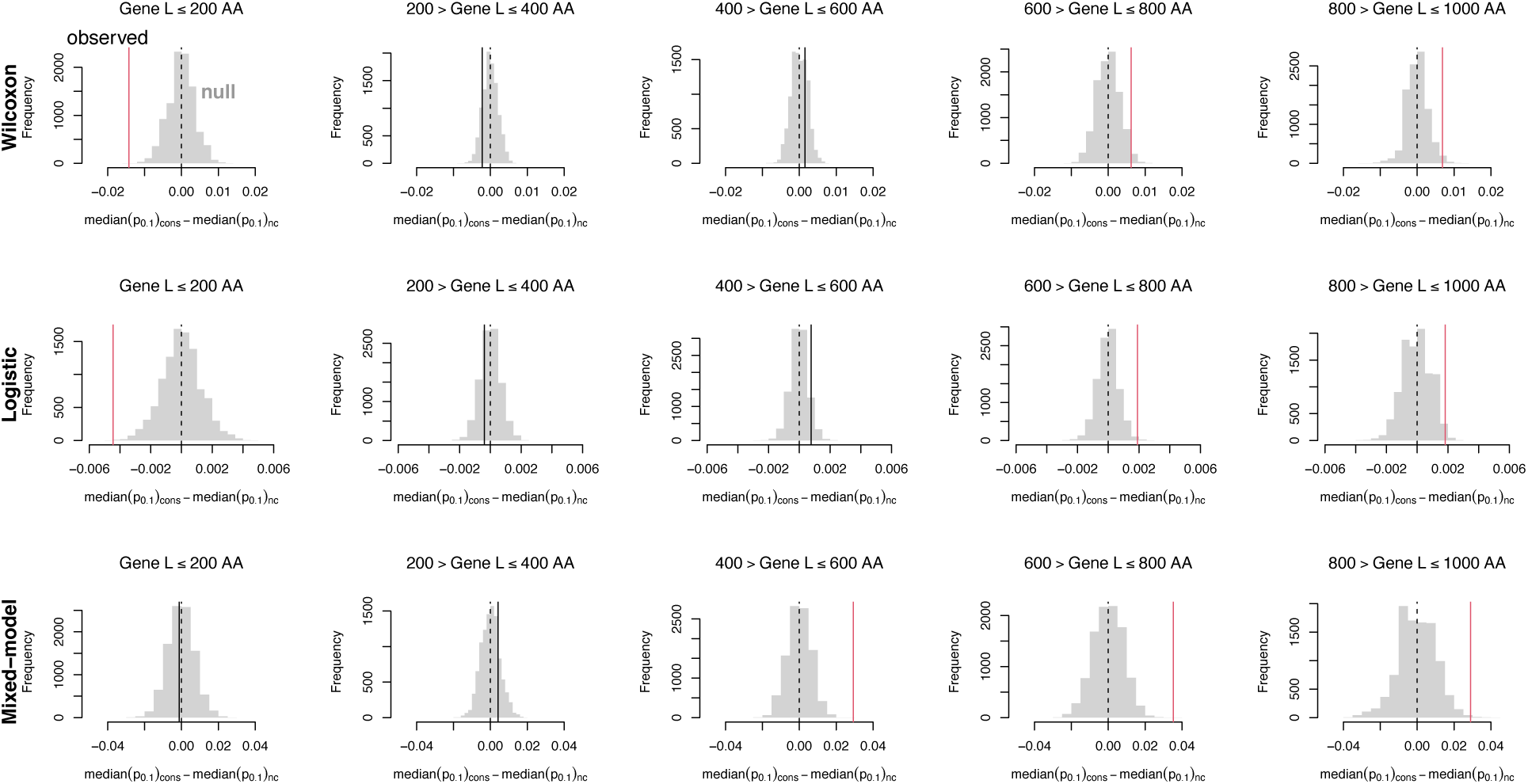
Comparing the observed difference in temporal allele shifts in conserved (‘cons’) versus non-conserved (‘nc’) genes of different lengths (AA = amino acid) for Eurasian samples. For each gene we calculated the 0.1 quantile p-value for SNPs in the gene, and then we compared the median of these values between conserved and non-conserved genes. When the observed is *less* than zero and the null, it indicates conserved genes show greater temporal allele frequency shifts (as for genes less than or equal to 200 AA in length using the Wilcoxon and logistic tests that do not correct for population structure). When the observed is *greater* than zero and the null, it indicates non-conserved genes show greater temporal allele frequency shifts (as for genes greater than 600 AA in length). The overall trends are similar between tests except for small genes, where the greater turnover in conserved small genes is only seen in tests that do not correct for population structure (Wilcoxon and logistic) while the mixed model turnover is not different for small, conserved genes compared to non-conserved. This discrepancy for small genes suggests that the high turnover for conserved small genes follows the genome-wide pattern for turnover, a turnover possibly due to drift being stronger due to their background selection. However, this hypothesis does not seem consistent with what is observed for longer conserved genes (less turnover than expected).

**Figure S5.**
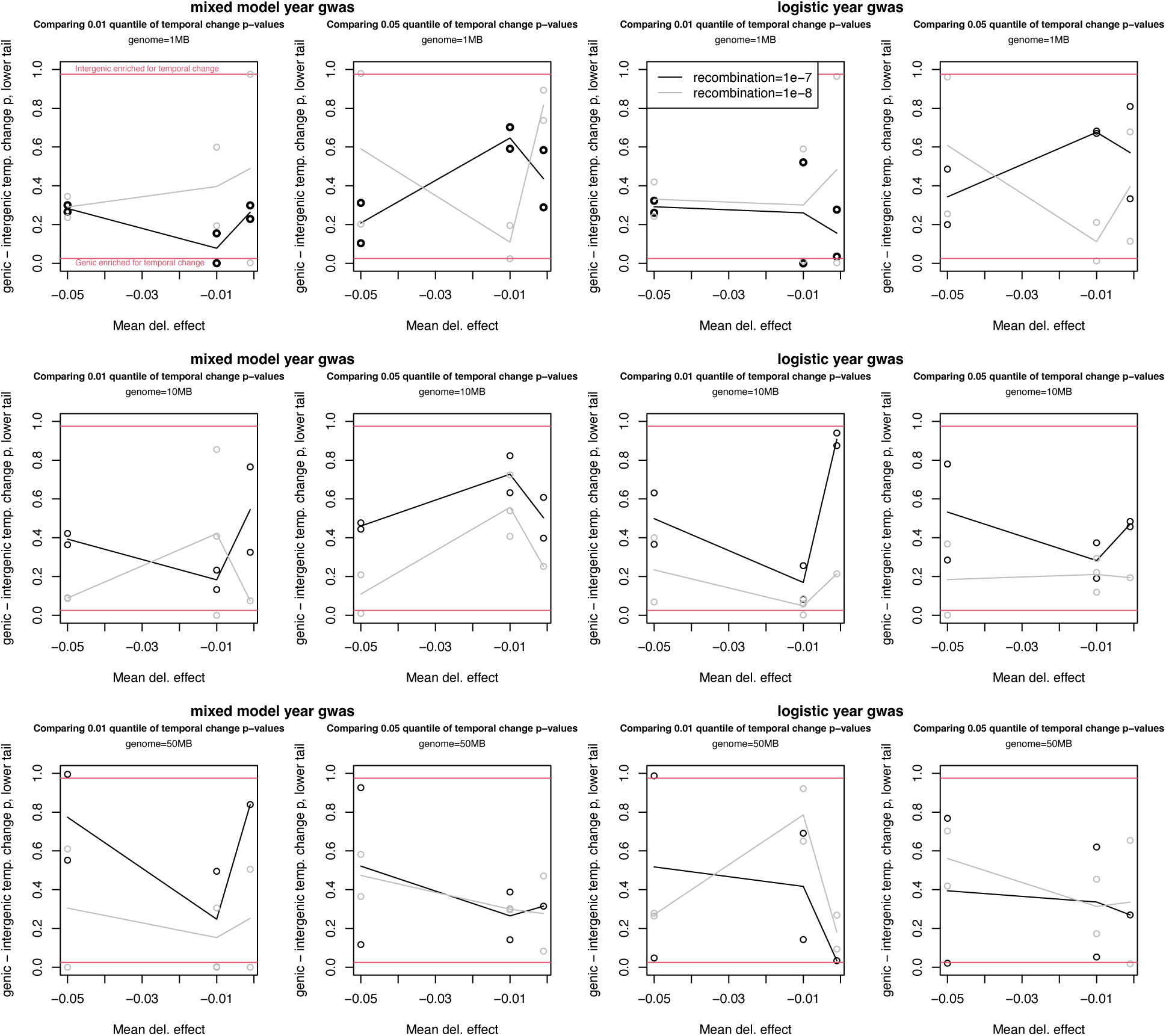
Results from simulations of purifying and background selection show no clear effect on the enrichment of genic SNPs for temporal allele frequency change. Purifying selection acts on a subset of coding SNPs and a subset of intronic SNPs, with neutrality for other SNPs. The y-axes show the lower tail of a permutation based null-model test, where the test statistic is the genic SNPs’ 0.01 p-value quantile minus the intergenic SNPs’ 0.01 quantile (1^st^ and 3^rd^ columns) or 0.05 quantile (2^nd^ and 4^th^ columns). Both linear mixed models testing year correlations with SNPs while accounting for kinship (1^st^ and 2^nd^ columns), or simple logistic year versus SNP allele (3^rd^ and 4^th^ columns) models are shown. Dots show individual simulation results, lines show means.

## Notes

### Competing Interest Statement

The authors have declared no competing interest.

### Summary of Updates

New versions of figures are included and a substantial text update is included.

